# Skin lipid chemistry influences host-microbiome-pathogen interactions in snake fungal disease (ophidiomycosis)

**DOI:** 10.1101/2025.11.04.686605

**Authors:** Kaitlyn M. Murphy, Jason W. Dallas, Mitra Ghotbi, Lluvia Vargas-Gastélum, Charlotte Van Moorleghem, Joshua L. Phillips, David Ludwig, Tingting Sun, Robert T. Mason, Joseph W. Spatafora, John H. Niedzwiecki, M. Rockwell Parker, Jeffrey D. Leblond, Justin M. Miller, Clay Stalzer, Ori Bergman, Melinda S. Chue Donahey, Marc G. Chevrette, Donald M. Walker

## Abstract

Within host-microbiome-pathogen systems, the host chemical microenvironment is often overlooked despite its inherent role in host physiology. We used a multifaceted experimental approach encompassing culture-dependent and independent methods, metagenomic and genomic data, and deep neural network modeling to assess the impact of host skin lipid chemistry and the bacterial microbiome on the growth of *Ophidiomyces ophidiicola* (ophidiomycosis, snake fungal disease). Results suggest that host skin lipid chemistry (e.g., oleic acid, squalene) and bacteria isolated from wild snake skins (e.g., *Chryseobacterium* sp. and *Stenotrophomonas maltophilia*) suppress *O. ophidiicola* growth. Notably, the *O. ophidiicola* genome contains biosynthetic gene clusters (BGCs) that encode metabolites that may suppress host lipid production, facilitating fungal pathogenicity. The contrastive deep neural network produced a near-perfect alignment of snake skin lipid and microbiome profiles for both individual snakes and disease states. BGCs from bacterial genomes isolated from snake skin overlap with metagenome profiles from wild snakes and correlate with disease state. We highlight antifungal activity found in the diverse lipid milieu of snake skin and bacterial-fungal interactions (BFIs) that structure the skin microbiome. Our results illustrate a strong relationship among a fungal pathogen, the microbiome, and host skin lipid chemistry.

## Introduction

Emerging fungal pathogens have highlighted the importance of acknowledging different spatial and temporal scales for mitigating disease outbreaks ^1,2^. The influence of abiotic and biotic factors on wildlife epidemics varies by pathogen distribution at local and regional levels ^3^, and by host life-history traits ^4^. However, the incorporation of biological scales (i.e., microbial to whole-organism) is often overlooked in identifying infection and development of disease ^5,6^. For example, chytridiomycosis is caused by the fungal pathogen *Batrachochytrium dendrobatidis* (*Bd*) and is responsible for the global decline of amphibian populations ^7^. Both landscape and local effects influence host infection and *Bd* prevalence ^8^, while diseased individuals have a distinct microbiome with microbe-derived metabolites that contribute to infection status ^9–11^. This shift in the microbiome is referred to as dysbiosis, a state marked by a reduction of beneficial microbial taxa ^12^, greater community dispersion ^13^, reduced resilience following pathogen clearance, ^14^ and can impair its benefits to the host. Due to the complex interactions between a host and pathogen, the integration of diverse data types, that approach questions of dysbiosis in multiple ways, are necessary to understand disease ecology and mitigate wildlife epidemics.

The community of microbes colonizing a host and the metabolites they produce are referred to as the microbiome^15^. The microbiome interacts with both the host and environmental factors and in part determines healthy or disease phenotypes ^16^. The microbiome acts as an extension of the immune system ^17,18^ to confer disease resistance through direct competition with pathogens, metabolite production, or stabilizing community structure to support cellular defense ^2,19^. Typically, studies of wildlife diseases focus on exploring the microbiome and metabolome in field settings, comparing diseased and healthy individuals to identify correlative links between microbial taxa and important phenotypes like disease state ^9,20–22^ and community resiliency ^9,14,20,21,23^. Culture-based studies have identified microbes that inhibit pathogens *in vitro* and provide evidence for putative function in disease resistance in free-ranging hosts ^24,25^. While these results provide insight into host relevant microbes, they fail to acknowledge a significant factor in the host-microbiome-pathogen system – the chemical composition of the host microenvironment and its influence on pathogenicity and disease.

The skin of all terrestrial vertebrates produces and maintains a rich mixture of both polar and nonpolar lipids that evolved to retard transcutaneous water loss ^26^. Squamate reptiles (i.e., lizards and snakes) harbor a complex lipid matrix ^27–29^. Functionally, snake skin broadly serves three major roles: i) to regulate permeability, ii) semiochemical communication ^30,31^ and iii) to mediate antimicrobial defenses ^32^. Defense is accomplished by many lipid components of the matrix, especially free fatty acids that create a physical barrier to microbial colonization ^33^ and exhibit antifungal activity ^34^. However, there are bidirectional interactions between the skin microbiome and the skin lipid composition, as the former modifies the latter, through changes in the microbial community and the metabolome, while lipid composition can also influence microbiome assembly^35^. Additionally, interspecific interactions occur within microbiomes, influencing the growth of community members, ^36^ and outcomes dependent on the state of the environment^37^. Therefore, the interplay between the skin microbiome and its environment could have downstream effects on the immune enhancing function of the microbiome. Tractable and manipulatable synthetic systems that utilize biologically relevant conditions have the potential to improve our fundamental understanding of non-model host-microbiome-pathogen systems. Generalizable trends in disease ecology may arise when results from synthetic systems are coupled with live animal studies and observations on the landscape.

*Ophidiomyces ophidiicola*, the causative agent of ophidiomycosis (common name: snake fungal disease [SFD]), has been identified around the world in both captive and wild snake populations ^38–40^. The fungus exhibits lipase and keratinase activity to successfully colonize snake skin ^39^. The pathogen drives dysbiosis in the skin microbiome, as demonstrated through both large-scale field and captive studies ^41^, likely via enzymatic alteration of the metabolic niche space on host skin. In this study, we characterize the interplay between *O. ophidiicola*, snake skin chemistry, and the skin microbiome of both captive and free-ranging snakes. We assess *O. ophidiicola* growth on prominent host skin compounds, characterize differences in bacterial functional profiles in the presence/absence of this pathogen, and use contrastive deep learning models to integrate two very different data domains, and link the microbiome to host skin chemistry profiles.

To explore the relationships among host skin composition, *O. ophidiicola*, and skin-associated bacteria, we employed a multifaceted experimental approach, including: i) the role of skin lipids in modulating *O. ophidiicola* growth, ii) how *O. ophidiicola* influences the snake skin microbiome structure and function and concomitant changes in skin chemical composition, iii) the direct and indirect effects of *O. ophidiicola* and resident skin bacteria on one another’s growth, and iv) a genomic analysis of *O. ophidiicola* and resident skin bacteria to assess putative mechanisms underlying *in vitro*, *in vivo,* and landscape level results. Analyses of culture-dependent and -independent methods suggest that host skin lipids, and certain bacterial isolates inhibit the growth of *O. ophidiicola*. However, the *O. ophidiicola* genome exhibits *in silico* evidence of gene clusters involved in the biosynthesis of metabolites that may suppress host lipid production, potentially promoting fungal colonization of snake skin. Our results highlight the importance of integrating host chemistry into studies of bacterial-fungal interactions (BFIs) in disease ecology and illustrate a strong relationship among a fungal pathogen, the host microbiome, and host skin chemistry.

## Results

### Snake skin lipids reduce O. ophidiicola growth

To assess how snake skin lipids influence the growth of *O. ophidiicola*, lipids were extracted from snake skin of five species (Supplementary Table 1) from Oregon, USA in a 1:1 hexane:dichloromethane mix over 24 hours. Lipids were then resuspended in solvent and divided into three technical replicates (n = 17). The lipid suspension was added to Sabouraud dextrose agar (SDA) and the remaining solvent was evaporated off before the medium was poured into a plate. Additionally, a solvent control (n = 6; solvent minus lipids) and negative control (n = 12; only SDA) were also prepared. Plates were inoculated with *O. ophidiicola* and incubated for 19 days at which growth was measured.

*Ophidiomyces ophidiicola* growth was significantly reduced on media containing the lipids of snake skins from all five species relative to controls (Fig. 1). Although a suppressive effect from the solvent control was observed (β = -185.8, *SE* = 43.7, *P* < 0.001), all inoculated plates containing snake skin lipids still differed from solvent controls (e.g., *Thamnophis sirtalis*: β = -575.8, *SE* = 66.8, *P* < 0.001; Fig. 1A). The species exhibiting the greatest suppression of *O. ophidiicola* on snake skin lipid media was *Pituophis catenifer* (β = -688.8, *SE* = 56.4, *P* < 0.001; Fig. 1A).

**Fig. 1.**
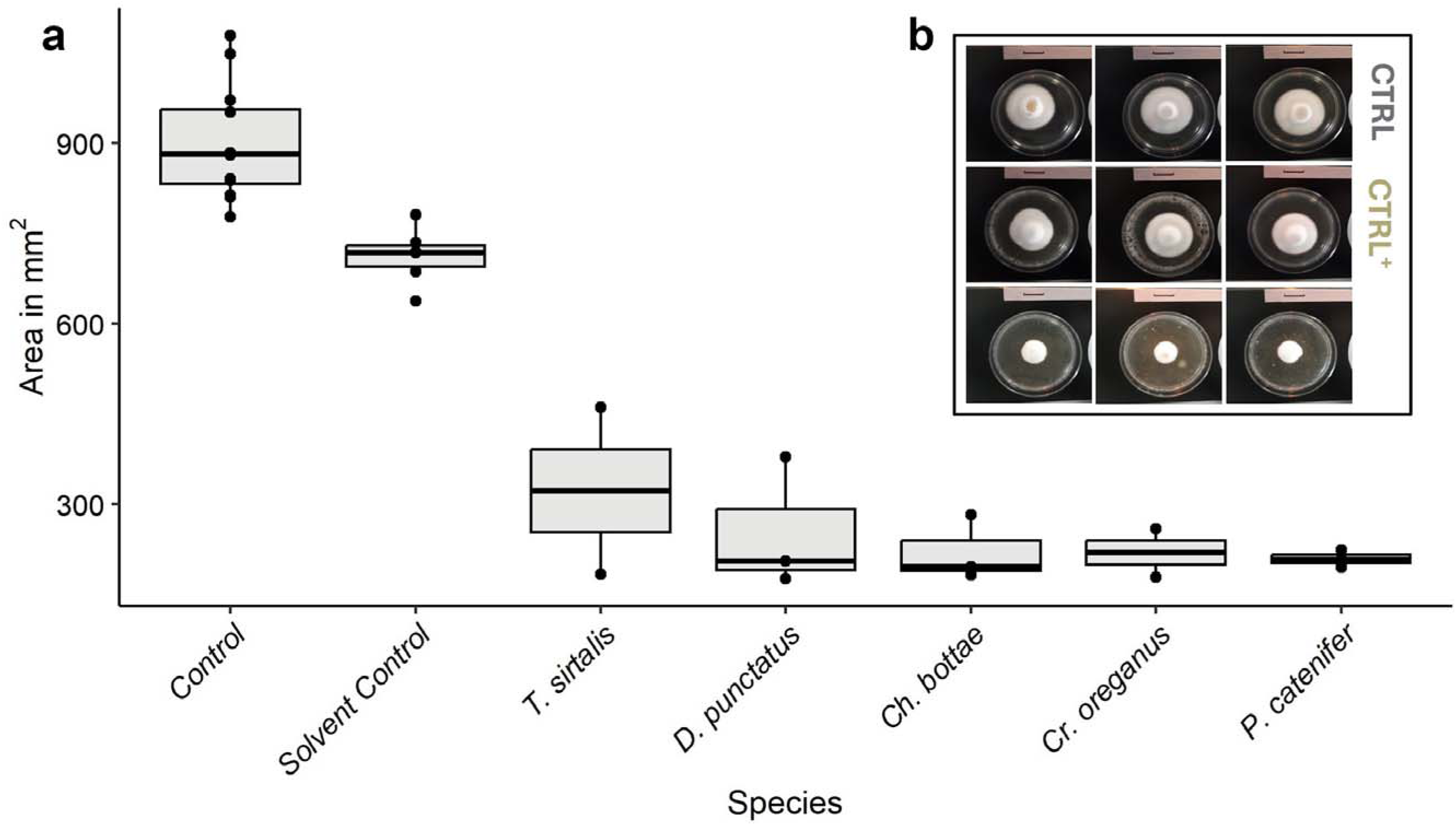
*Ophidiomyces ophidiicola* growth on media mixed with wild snake skin lipids. a) The area of *O. ophidiicola* growth (mm^2^) after 19 days with extracted lipids from 5 species of snakes and their respective controls (n = 31). All values are statistically significant (P ≤ 0.001) from controls. Center lines within box plots represent the median and the boxes denote the interquartile range, with whiskers representing 1.5× the upper or lower quartile. b) Differential growth patterns of *O. ophidiicola* grown in replicate on control, solvent control, and snake skin lipid media.

### O. ophidiicola growth varies across skin carbon source and concentration

Keratin, cholesterol, oleic acid, and squalene represent some of the most abundant snake skin compounds^28^ and were thus used to test for *O. ophidiicola* growth facilitation or inhibition in carbon minimal media at five concentrations (1, 10, 100, 1,000, 10,000 ppm), along with M9 carbon neutral medium as a control. Plates (n=5 replicates per treatment or control) were inoculated with *O. ophidiicola* and incubated at 25 °C for 22 days. The diameter of mycelial growth was measured on the final day. Additionally, we tested for an interactive effect of keratin and each lipid listed above using 10,000 ppm media under identical growth conditions.

Linear mixed effect modeling and *post-hoc* pairwise comparisons indicated that *O. ophidiicola* grows statistically differently on all four minimal media with numerous significant interactions between medium and concentration (n = 97, *P* < 0.001; Supplementary Table 2 – 4). Keratin minimal media supported the fastest growth of *O. ophidiicola* relative to the other media (β = 62.1 mm, *SE* = 0.9, *P* < 0.001; Fig. 2A, B), but growth rates decreased as keratin concentration was reduced (Supplementary Table 4). Cholesterol was the only carbon source that largely exhibited similar growth across all concentrations, with the only pairwise differences being between 10,000 ppm and 100 ppm (β = -5.0 mm, SE = 1.3, P = 0.02) and 1,000 and 100 ppm (β = -7 mm, *SE* = 1.3, *P* < 0.001; Supplementary Table 3), and 100 ppm being the only concentration different from controls (n = 42; β = 6.6 mm, *SE* = 1.9, *P* = 0.001; Supplementary Table 4). Both oleic acid and squalene suppressed *O. ophidiicola* at high concentrations (10,000 and 1,000 ppm), with the latter completely inhibiting fungal growth, but the suppressive effects were lost at lower concentrations as 10 and 1 ppm were similar to controls (n = 42, 40; β = -1 – - 2.3 mm, *SE* = 1.4 – 1.5 mm, *P* > 0.09; Supplementary Table 4).

**Fig. 2.**
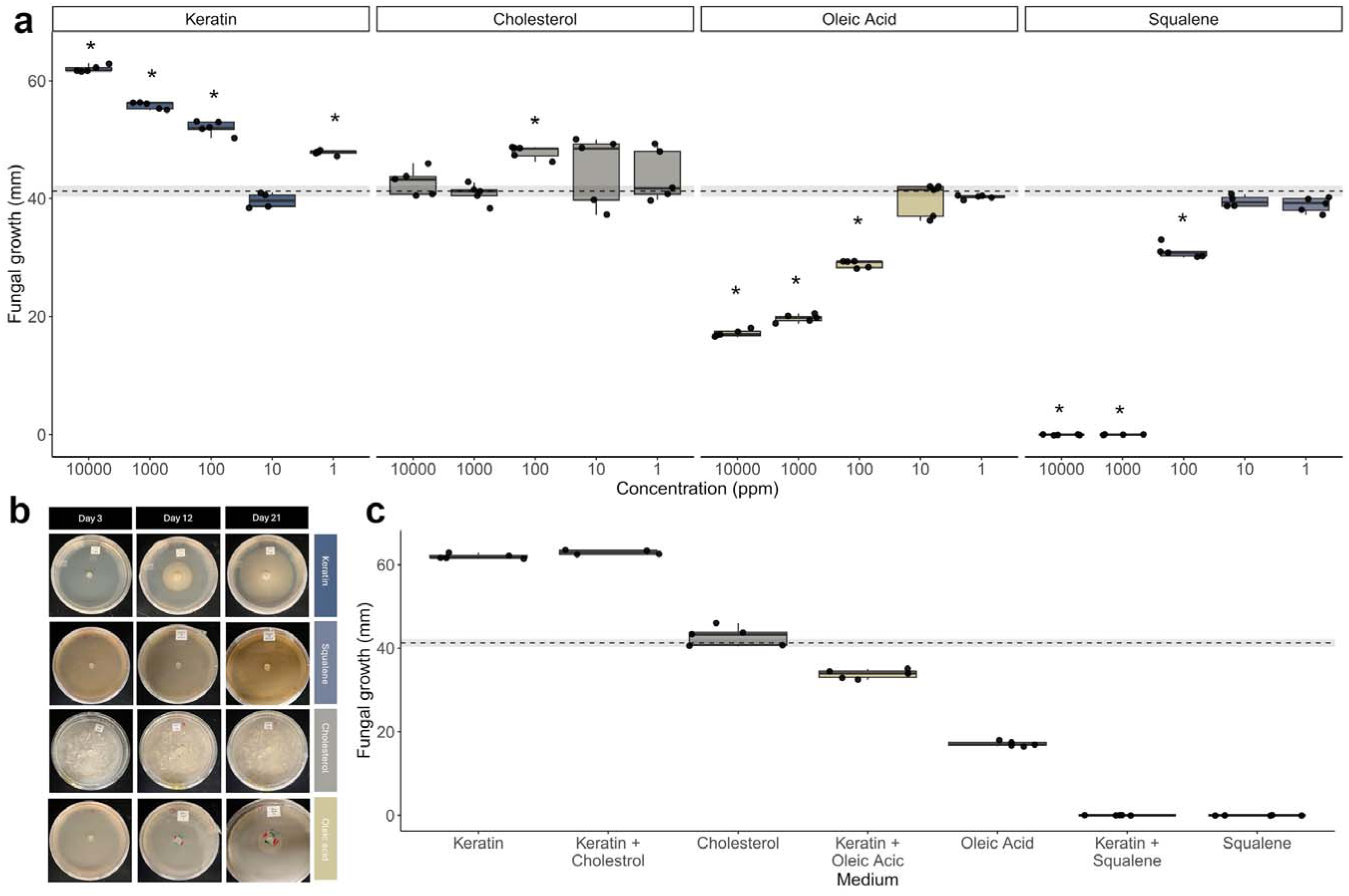
*Ophidiomyces ophidiicola* growth on minimal media sourced from common compounds found in snake skin. a) Fungal growth on plates with single carbon sources of keratin, cholesterol, oleic acid, or squalene. The concentrations of the carbon source are shown in descending order along the x-axis. The dashed line and shaded area represents the mean and ± 1 standard error *O. ophidiicola* growth on the M9 minimal media control. Asterisks above the box plots represent significant differences in *O. ophidiicola* growth compared to the M9 minimal media control (see Supplementary Table 4). b) Representative images of *O. ophidiicola* growth on 10,000 ppm minimal media over time. c) Interaction effects between keratin and each lipid at 10,000 pm concentration of each carbon source. The dashed line and shaded area represents the mean and ± 1 standard error *O. ophidiicola* growth on the M9 minimal media control. For both a and c, center lines within box plots represent the median and the boxes denote the interquartile range, with whiskers representing 1.5× the upper or lower quartile.

When keratin and lipids were combined in mixed carbon source media, fungal growth was restricted in plates containing squalene (n = 34; β = -62.1, *SE* = 0.4, *P* < 0.001) and oleic acid (β = -28.3, *SE* = 0.4, *P* < 0.001) when compared to keratin plates (Fig. 2C; Supplementary Table 5). Addition of cholesterol resulted in a minor increase in *O. ophidiicola* growth relative to keratin plates, but this difference only approached significance (β = 1.0, *SE* = 0.4, *P* = 0.052).

### Contrastive deep neural network modeling of the skin microbiome and host lipid chemistry

To explore the interplay between the microbiome and skin lipid composition in the presence of *O. ophidiicola*, juvenile Common Water Snakes (*Nerodia sipedon;* n = 22) were either inoculated with *O. ophidiicola* or a sham control and monitored over 80 days, or until an individual experienced mortality (see Romer et al. ^42^ for detailed methods). A skin microbiome sample was collected weekly and the 16S-V4 rRNA marker sequenced and clustered into operational taxonomic units (OTUs) at 97% similarity. Upon the conclusion of the experiment, all snakes were euthanized and preserved in formalin. Carcasses were rinsed of formalin with DI water for one minute before being submerged in a 1:1 hexane:dichloromethane mix for 24 hours to extract skin lipids with the head and cloaca kept out of the beaker in order to avoid possible extraction of internal lipids. The extracted lipids were then analyzed using gas chromatography-mass spectrometry (GC/MS).

Contrastive deep learning methods were used to compare patterns in the skin microbiome to host lipid chemistry from the live animal study. Amplicon samples were processed using a modified version of the DNABERT model ^43^ that was pre-trained for 100,000 steps on the SILVA v138.1 16S dataset ^44^. GC/MS samples were processed using a standard transformer model based on vision transformer ^45^. The retention times were mapped directly to sinusoidal position encodings and the TiC signals were normalized. The amplicon and GC/MS models were trained in tandem using contrastive pre-training for 30,000 steps. A batch consisted of six paired amplicon and GC/MS samples. The models were trained following the contrastive pre-training objective as defined in Radford et al ^46^. In order to align the latent spaces, we shared the weights of the final linear projections and added an additional mean-squared error loss between the paired embeddings.

Using contrastive pretraining methods, we integrated two deep learning models to link skin chemistry to dysbiosis of the microbiome (Fig. 3). By contrastively training the amplicon and GC/MS encoders on paired sample data, the models were able to learn shared latent space representations aligning sample embeddings from the two different data domains (Fig. 3A). The models effectively paired skin chemistry and microbiome profiles between samples and organized the latent space representing *O. ophidiicola* presence in an unsupervised manner.

**Fig. 3.**
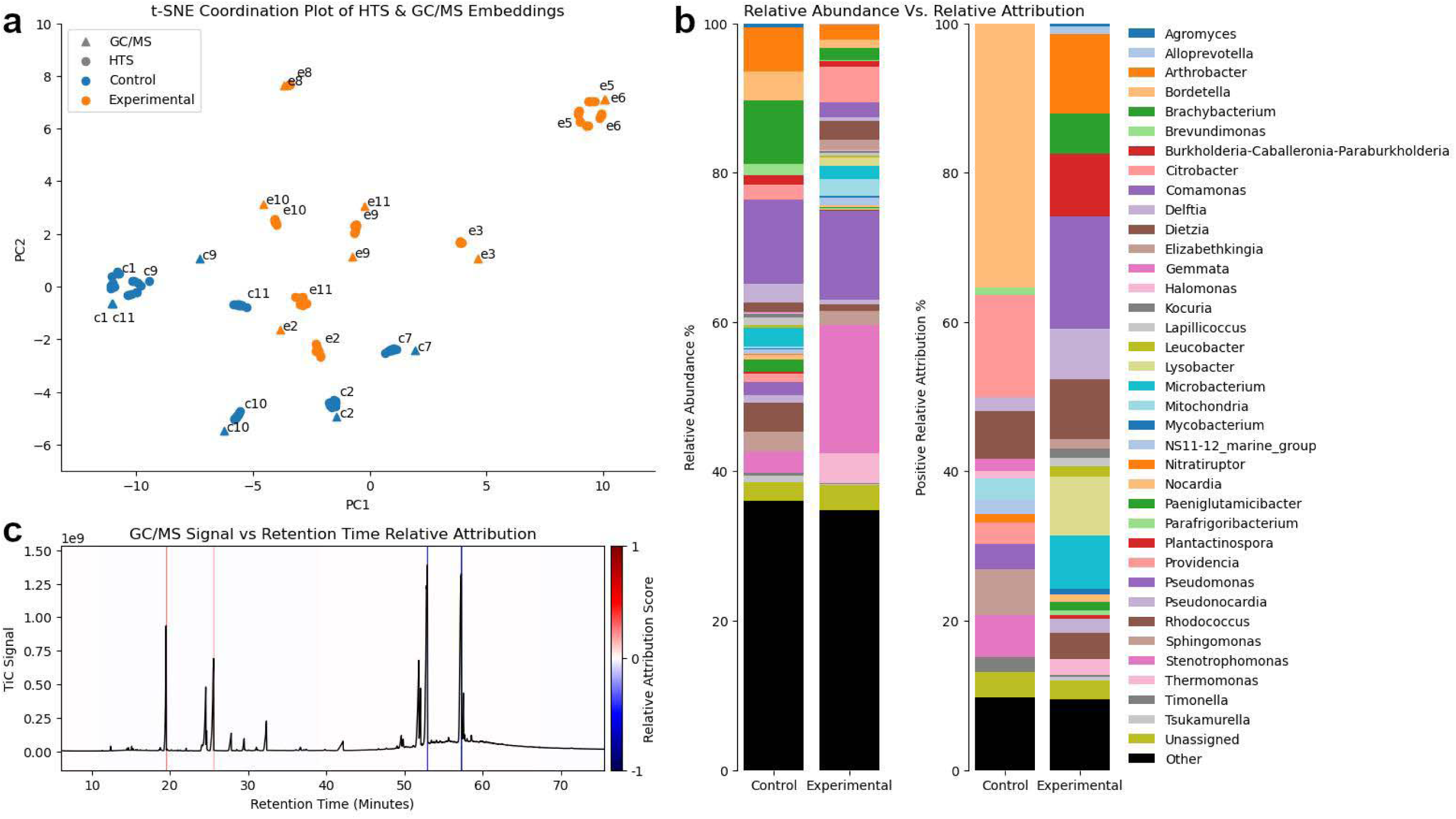
Examining the relationship between the snake skin microbiome and its lipid profile. a) The resulting t-SNE plot of the sample-level embeddings produced by the contrastively-trained amplicon and GC/MS encoders. The GC/MS points represent the embeddings produced by the GC/MS encoder given the entire profile. The amplicon points represent subsample embeddings resulting from the amplicon encoder, where each sample is subsampled independently 10 times, and each subsample consists of 1,000 sequences drawn randomly with replacement. b) The relative abundance and corresponding positive relative attribution scores resulting from the amplicon encoder. The top 90% explained genera were drawn from the total positive attribution scores and shown. c) The GC/MS profile of e2 with a banded background representing the relative attribution produced by the GC/MS encoder at the given time points. Positive attribution indicates that the signals at the corresponding times drive class prediction, while negative attribution scores hinder prediction.

We employed self-attention attribution from the transformer architecture to explain the encoder embeddings. We fine-tuned each encoder independently to predict the presence of *O. ophidiicola*, holding all parameters constant, except for the final linear projection. Self-attention attribution was computed for each amplicon and GC/MS sample using the fine-tuned encoders to capture bacterial genera explaining 90% of the attribution score from the amplicon encoder (Fig. 3B). The relative attribution differs significantly from the relative abundance scores indicating that the model is leveraging more complex interactions within the microbiome to make predictions as opposed to those based on relative abundance shifts. For the GC/MS profiles, the attribution scores for each snake were obtained and plotted against the original profiles to identify peaks representative of *O. ophidiicola* state (Fig. 3C; Supplementary Fig. 1).

### Direct Interactions between Snake Skin Bacteria and Ophidiomyces Inhibit Fungal Growth

To examine BFIs between *O. ophidiicola* and snake skin bacteria, a direct bacterial-fungal interaction (DBFI) experimental setup was implemented. To begin, six bacterial isolates (Supplementary Table 6) were streaked onto 10,000 ppm keratin minimal medium and grown at 25 °C for 48 hrs while *O. ophidiicola* was grown on M9 minimal medium plates at 25 °C for 12 days. The following treatments and controls were set up in triplicate in a fully factorial design including: 1) *O. ophidiicola* with/without a bacterial isolate on a 10,000 ppm keratin carbon source, 2) *O. ophidiicola* with/without a bacterial isolate on M9 (minus host carbon source), and 3) bacterial isolate (no *O. ophidiicola*) on 10,000 ppm keratin carbon source or on M9 (minus host carbon source) in 12-well plates for a total of 84 samples. A ten-fold serial dilution of bacterial isolates was carried out, optical density 620 nm measured, and the dilution reading 0.15 used to inoculate each the experimental replicate containing bacteria ^47^. Plates were incubated at 25 °C and passaged every 24 hrs over a week, by aspirating broth with bacterial growth, and replenishment with fresh broth into each well. At each passage event, bacterial and fungal growth were simultaneously quantified by measuring the OD620 nm for bacteria and *O. ophidiicola* radius of growth.

When six single bacterial isolates and *O. ophidiicola* were grown in co-culture, a range of interactions were observed in terms of bacterial and fungal growth (Fig. 4; Supplementary Table 6). Some bacteria, including *Stenotrophomonas maltophilia*, grew more on keratin medium regardless of *O. ophidiicola* presence (*F*_1,8_ = 755, *P* < 0.001), while fungal growth was reduced in the presence of this bacterium (β = -0.12, *SE* = 0.02, *P* < 0.001). *Chryseobacterium* sp. growth, while greater on the keratin medium, was enhanced in the presence of the fungus (*F*_1,8_ = 5.62, *P* = 0.045). However, in the presence of Chryseobacterium sp., *O. ophidiicola* was restricted in growth only on keratin media (β = -0.11, *SE* = 0.15, *P* < 0.001). Both bacterial and fungal growth on M9 did not differ based on presence/absence of a paired member in co-culture suggesting that observed BFIs are driven by keratin availability. Five of the six bacterial species show antagonistic effects against *O. ophidiicola,* whereas *Acinetobacter* sp. does not interact with *O. ophidiicola* (*F*_1,8_ = 4.11, *P* = 0.339). For a full description of the effects of co-culturing on fungal and bacterial growth, see Fig. 4 and Supplementary Table 6.

**Fig. 4.**
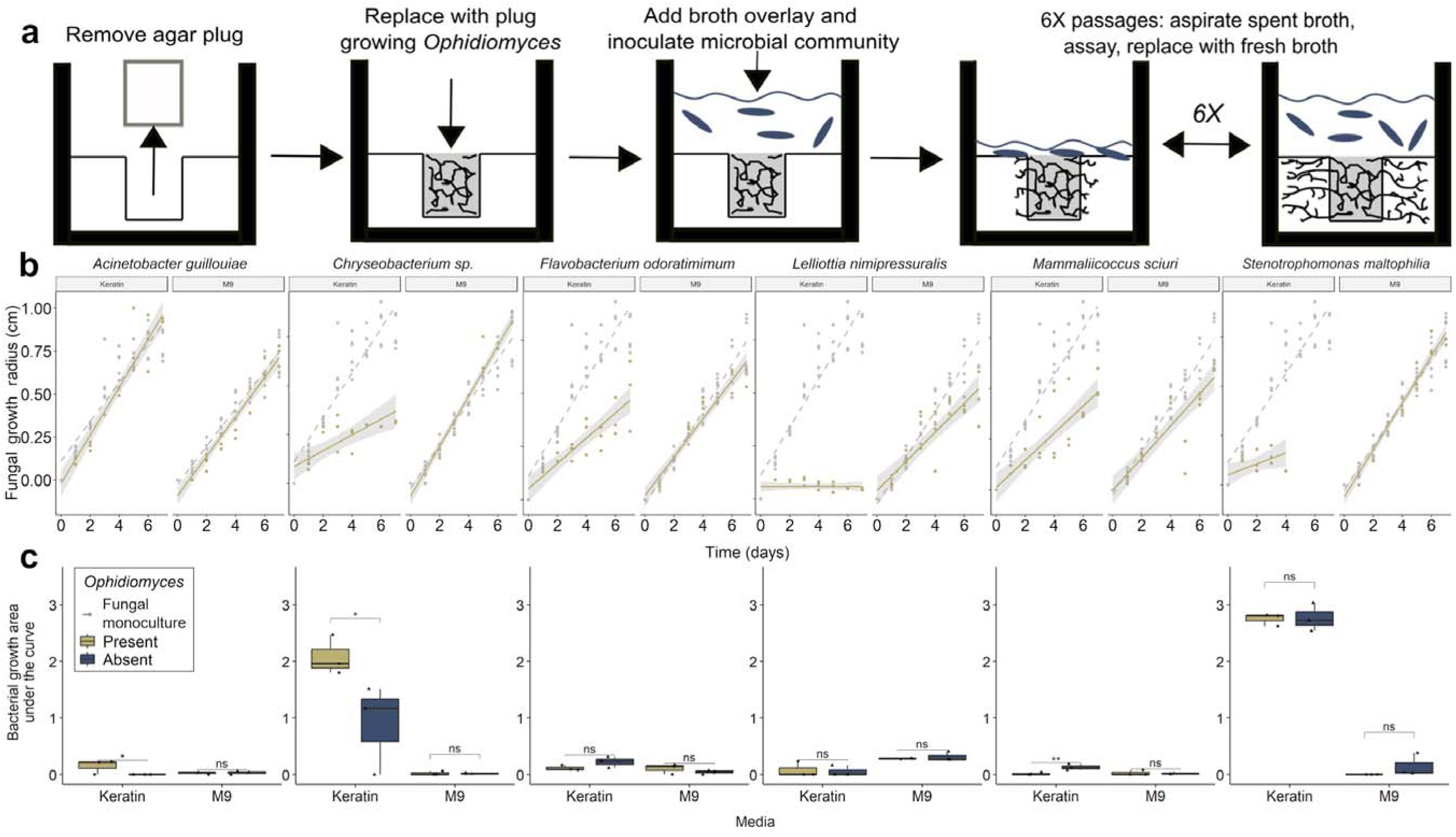
Growth of bacteria and *Ophidiomyces ophidiicola* in a direct bacterial-fungal interaction (BDFI) assay. a) Experimental design of the DBFI method. In this study, experiments were performed in 12-well plates and inoculated with *O. ophidiicola*. Plates were then overlaid with a monoculture broth and passaged 6X, once every 24 hours, to maintain nutrient availability. At each passaging event, both *O. ophidiicola* and bacterial growth were recorded. b) Growth of *O. ophidiicola* was differently affected by the overlaying bacterial monoculture relative to an *O. ophidiicola* monoculture. c) Bacterial growth was compared between wells with and without growing *O. ophidiicola*. Center lines within box plots represent the median and the boxes denote the interquartile range, with whiskers representing 1.5× the upper or lower quartile.

### Fungal exometabolites have a variable response on bacterial growth

Keratin media metabolized by *O. ophidiicola* (fungal spent media) was used to explore the indirect effects of a fungal modified environment on bacterial growth, as bacterial communities are known to respond directly to fungal competition for a nutrient source or fungal-derived exometabolites ^48^*. Ophidiomyces ophidiicola* mycelia were added to three 2.8 L shaker flasks containing 1 L of 10,000 ppm keratin and flasks were incubated at 25 °C at 120 rpm. To create a gradient of resource usage or metabolite production by *O. ophidiicola* (Fig. 5A), a single shaker flask from each carbon source at days 5, 9, and 13 was filtered through autoclaved cheesecloth to remove mycelia and collect the spent media (SM). Glycerol stocks of bacteria isolated from snakes (n = 18) were inoculated in triplicate as monocultures into 10,000 ppm keratin minimal media (positive control), M9 minimal salts (negative control), 5, 9 or 13 day SM (n = 285). All experiments were conducted in 96-well flat bottom plates which were incubated at 25 °C and absorbance reading values at 620 nm were recorded each hour for 50 hrs to measure bacterial growth.

**Fig. 5.**
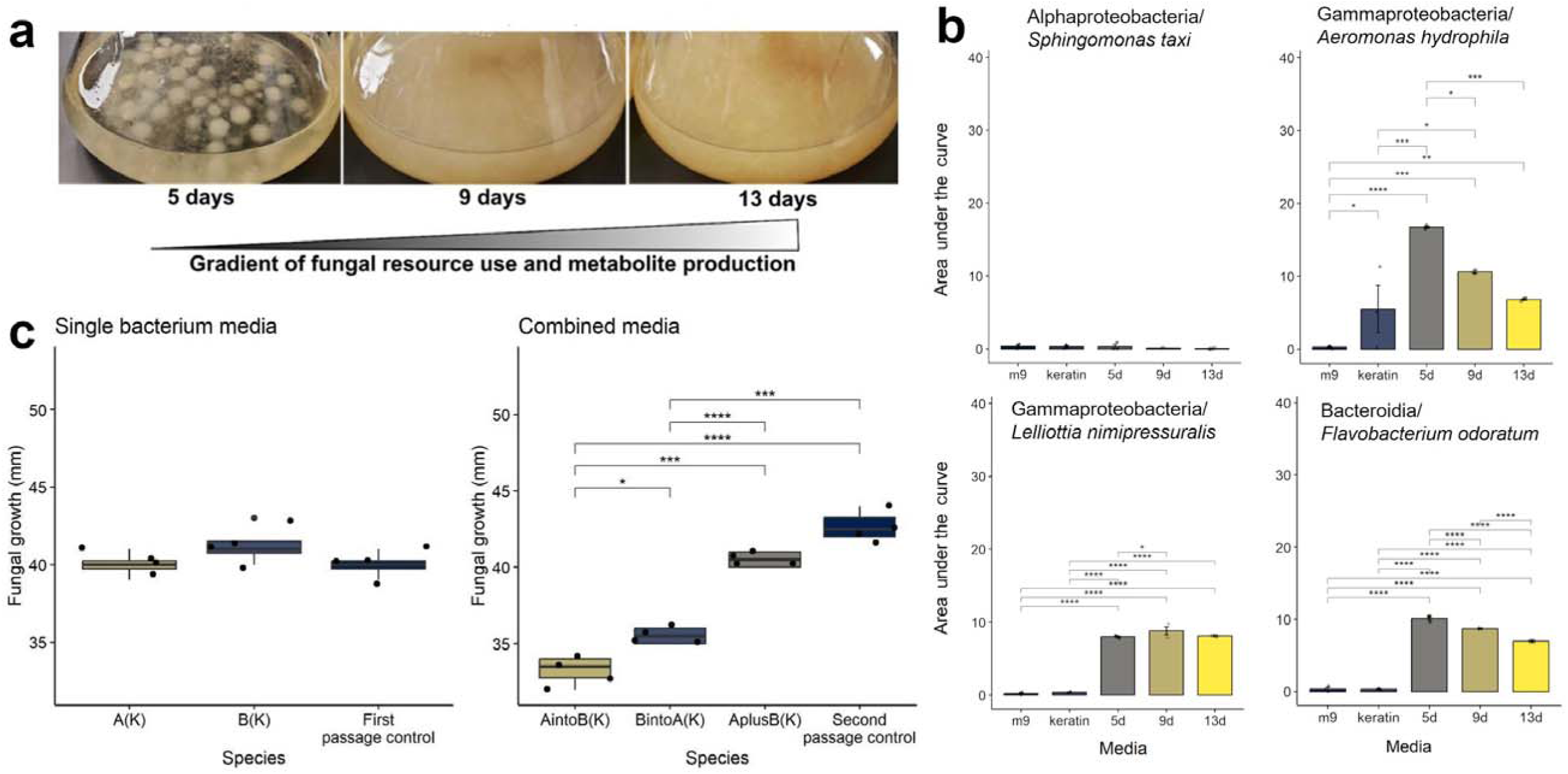
The growth of bacteria on *Ophidiomyces ophidiicola* metabolized keratin and vice versa. a) Representative images of fungal growth on 10,000 ppm keratin media over time. b) The effect of *O. ophidiicola* SM at 0 (“keratin”), 5, 9, or 13 days of fungal metabolism of a 10,000 ppm keratin medium for four different classes of bacteria. The four trends are general representatives of distinct trends of bacterial growth on fungal SM. Error bars denote ± 1 standard error. c) The effect of two bacterial strains on *O. ophidiicola* growth using bacterial spent media (SM) to test for monoculture interactions (single bacterium medium) between *O. ophidiicola* and *Chryseobacterium* sp. “A” or *Stenotrophomonas maltophilia* “B” or testing for cross-feeding (“A into B” or “B into A”) and additive interactions (“A plus B”) of *Chryseobacterium* sp. and *S. maltophilia*. These experiments were completed on a 10,000 ppm keratin minimal medium with M9 (carbon negative) control. Center lines within box plots represent the median and the boxes denote the interquartile range, with whiskers representing 1.5× the upper or lower quartile. Statistically significant pairwise differences via a Tukey HSD are indicated by: p < 0.05, *; p < 0.01, **; p < 0.001, ***; and p < 0.0001, ****.

Four broad patterns of bacterial growth in response to fungal SM were observed (Fig. 5B; Supplementary Fig. 2; Supplementary Table 7): 1) no bacterial growth across all media types (3/18 bacterial isolates), 2) linear, concentration-dependent relationship between SM type and bacterial growth, while also showing bacterial growth on the keratin control (3/18 bacterial isolates), 3) linear, concentration-dependent relationship that is completely reliant on fungal SM type as the bacteria do not grow solely on the keratin control (2/18 bacterial isolates), and 4) a non-linear concentration dependent relationship that required *O. ophidiicola* metabolites to grow (10/18 bacterial isolates). For example, *Sphingomonas taxi* exhibited no interactions between media types and bacterial growth (*P* > 0.05; Fig. 5B). However, *Aeromonas hydrophila* had the greatest growth on 5-day SM relative to other SM and the keratin and M9 controls (*P* < 0.05; Fig. 5B). Other bacterial taxa like *Lelliottia nimipressuralis* showed similar trends, but relied on *O. ophidiicola* metabolites to grow, as no statistical differences were observed between the keratin and M9 controls (*P* > 0.05; Fig. 5B).

### Bacterial Cross Feeding Suppresses Fungal Growth

To determine if *O. ophidiicola* responds to a bacterial modified skin environment, we performed the inverse of the previous experiment by growing *O. ophidiicola* on SM from two strains of bacteria that were identified using a neural network as important contributors to disease state ^41^. We also explored the possibility of bacterial cross-feeding on *O. ophidiicola* growth using the same two strains of bacteria*. Chryseobacterium* sp. (isolate BR1.10) and *S. maltophilia* (isolate TR087-6.4s) were grown on 10,000 ppm keratin or M9 control broth and incubated at 25°C while shaking. After 24 hrs, broth cultures were passed through a 0.22 uM filter to remove bacterial cells and collect the SM. The SM was partitioned into two aliquots including 60 mL for *O. ophidiicola* growth assays and 40 mL for bacterial cross-feeding experiments. *Chryseobacterium* sp. or *S. maltophilia* were inoculated into 40 mL of the opposite strain’s SM in a 75 mL flask then incubated at 25 °C while shaking for 24 hours. The liquid SM was added in an equal volume to a molten 2X agar medium, under constant mixing, and split into four replicate Petri plates per treatment or control to test for 1) the effect of independent bacterial strains on *O. ophidiicola* growth, 2) additive effects by combining the SM of each bacterial strain, and 3) cross-feeding of the SM from both bacterial strains and the impact on *O. ophidiicola* growth. Fungal growth on bacterial SM was measured after 14 days (n = 28).

When grown on SM from individual bacterial taxa (*Chryseobacterium* sp. or *S. maltophilia*), no difference in growth was observed (*P* > 0.05; Fig. 5C). However, when bacterial SM was cross-fed between the two bacterial isolates, fungal growth was observed to decrease when *Chryseobacterium* sp. SM was fed to *S. maltophilia* (β = -4.50, *SE* = 1.12, *P* < 0.001) and vice versa (β = -3.38, *SE* = 1.12, *P* = 0.006). Interestingly, no additive effect was observed (β = - 0.63, *SE* = 1.12, *P* = 0.58) when *O. ophidiicola* was grown on equal volumes of SM produced independently by *Chryseobacterium* sp. and *S. maltophilia*.

### Genomic predictions of interactions between skin bacteria and O. ophidiicola

To characterize the functional potential of bacterial communities during their interaction with *O. ophidiicola*, we assembled 13 bacterial metagenomes (MAGs) from 120 wild caught watersnakes (Fig. 6; Supplementary Fig. 3). Six MAGs were specific to *O. ophidiicola* positive and seven in disease negative snakes. AntiSMASH predicted different biosynthetic gene clusters (BGCs) profiles for disease-positive or -negative snakes. In disease-positive snakes, we found a 66% match to an NI-siderophore producing aerobactin and 100% identity to an NRP-metallophore producing enterobactin, while in negative snakes, we found a 75% match to the polyketide producing flexirubin, 33% and 66% identity to the NI-siderophores producing fulvivirgamide and desferrioxamine (respectively), and 100% match to an ε-Poly-L-lysine NRP.

**Fig. 6:**
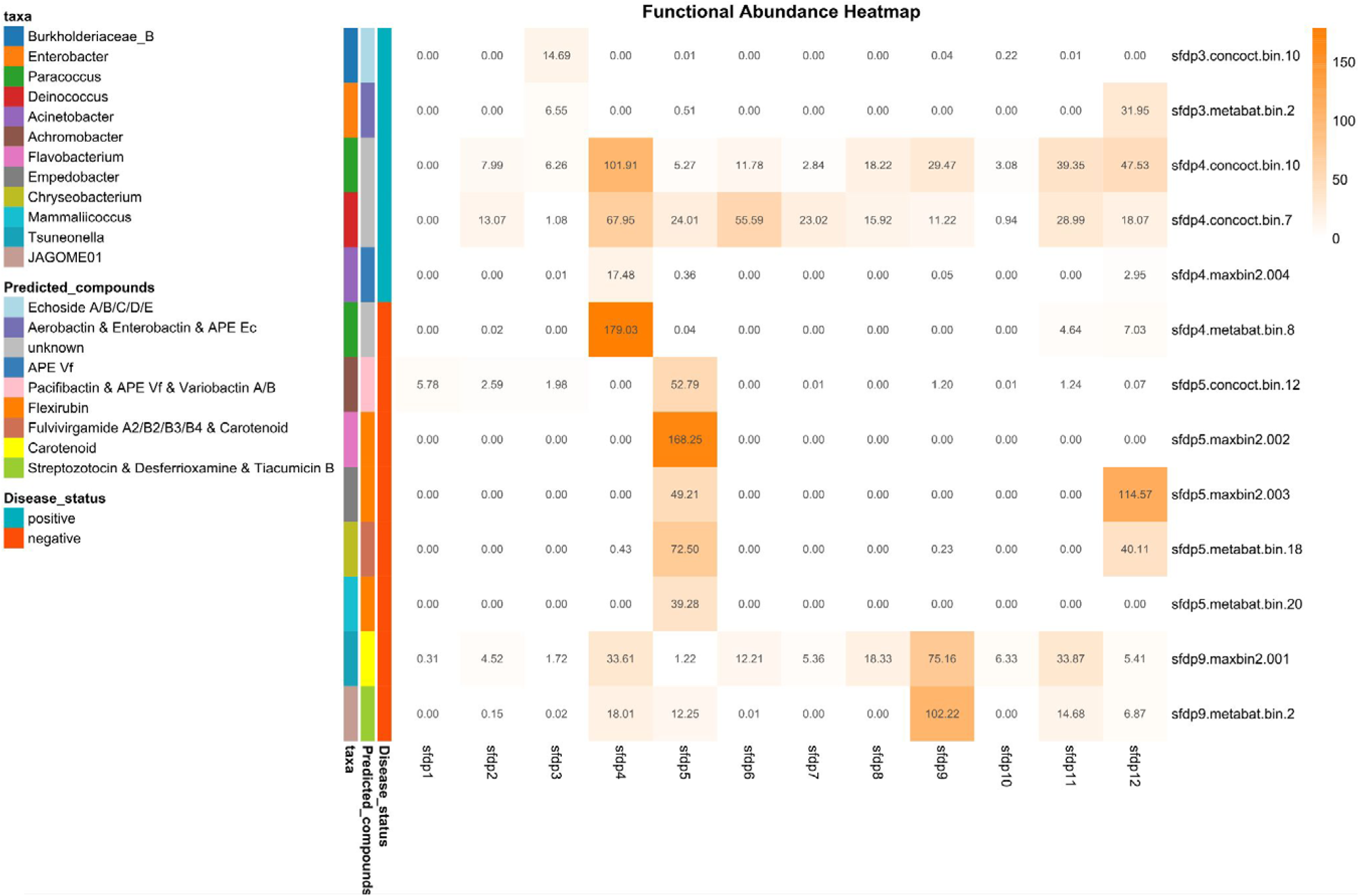
Heatmap illustrating the relative abundance of metagenome-assembled bacteria and their gene copies across different disease statuses. Metabolite types, extracted from the antiSMASH database, are annotated and color-coded on the left side of the heatmap. The dendrogram, constructed using Euclidean distance, clusters the taxa based on similarity. The heatmap is row-scaled to normalize values for comparison.

An analysis of BGCs across 26 living bacterial isolates revealed distinct patterns in both BGC richness and rediscovery of known compounds (Supplementary Table 8). A total of 14 BGCs in 12 isolates were identified by antiSMASH as siderophores suggesting iron or other metals are an important mediator of fitness in this environment. *Streptomyces* sp. (EKS17.29) had the highest number of BGCs (74 total, 50 on contig edges) while *Rhodococcus erythropolis* (EKS20.4) and *Chryseobacterium* sp. (BR1.10) had greater than 30 BGCs. *Streptomyces* sp. (EKS17.29) harbored desferrioxamin B among its rediscovered compounds, along with geosmin and other secondary metabolites. Additionally, a *Paenarthrobacter* sp. (EKS20.8) had the BGC for desferrioxamine E, another siderophore with iron chelating activity.

Secondary metabolites are believed to play a crucial role in the infection process by pathogenic fungi ^49^. AntiSMASH analysis on the *O. ophidiicola* genome resulted in the identification of a total of 25 putative secondary metabolite BGCs. Of these, 15 BGCs were classified as belonging to the categories of non-ribosomal peptide synthetases, polyketide synthases, and terpenes (Supplementary Table 9). In the *O. ophiidicola* genome, the gene cluster involved in the biosynthesis of Squalestatin S1 has several hypothetical proteins including the core enzyme farnesyl-diphosphate farnesyl/transferase and a DnaJ domain protein. Additional BGCs identified may be involved in the colonization process and are known to interact negatively with other members of the skin microbiome: these include griseosulfin (antifungal activity), chaetolivacine A (antimicrobial activity), and metachelin C, a siderophore important for acquisition of iron or other metals ^50^. Multiple proteases implicated in virulence including subtilisin-like serine proteases (sub-4, sub5 and sub9) along with leucine aminopeptidase (Lap1 and Lap2), extracellular metalloproteinase, and dipeptidyl-peptidase were annotated.

## Discussion

Studies spanning multiple scales in disease ecology allow for explicit hypothesis tests, cross validation of observed trends, and establishment of fundamental rules governing the development of disease. We used the SFD system to improve understanding of the role of host chemistry in disease ecology and BFIs contributing to dysbiosis. We found that fungal growth is negatively affected by snake skin lipids even if supplemented with growth enhancing keratin. Deep neural networks linked the microbiome to host chemistry and disease state. More specifically, the models accurately paired skin lipid profiles of individual snakes with their respective microbiomes (and vice versa), clustered snakes by disease state, and used attention attribution to identify bacterial taxa and lipid compounds explaining disease state. We also show that when bacteria are grown in co-culture with *O. ophidiicola* and on fungal SM, cooperative, competitive, and cross-feeding interactions were observed. We identified biosynthetic gene clusters (BGCs) in the genomes of bacterial skin isolates and *O. ophidiicola* that we predict govern BFIs in diseased snakes and showed commonalities with BGCs from bacterial metagenomes on the landscape. Our approach used a multifaceted dataset across multiple biological scales to improve our understanding of disease ecology, BFIs, and role of skin chemistry in fungal pathogen induced dysbiosis.

The skin of all terrestrial tetrapods produce a complex lipid matrix with many functions, including enhancement of a pathogen barrier ^51^. When *O. ophidiicola* was grown on media containing snake skin lipids, fungal growth was significantly reduced (Fig. 1). Despite species-specific differences in lipid composition in snakes ^52^, the complex lipid makeup of snake skin equally restricted *O. ophidiicola* growth. Allender et al. ^39^ found that *O. ophidiicola* exhibits lipase activity, indicating the capability of *O. ophidiicola* to metabolize complex lipids and interesting interplay between fungal growth suppression and enzymatic activity. To isolate the effects of specific host lipids on *O. ophidiicola*, we focused on cholesterol, oleic acid, and squalene, which are dominant, ubiquitous components of the snake skin lipid matrix ^30,52–54^. Both oleic acid and squalene had strong suppressive effects on *O. ophidiicola* at the three highest concentrations even in the presence of growth promoting keratin. Squalene is a ubiquitous component of terrestrial vertebrate skin lipids, and studies on human skin dysbiosis with fungal pathologies use drugs (e.g., terbinafine) that enhance squalene persistence in skin to suppress fungal growth ^55^. The presence of squalestatin S1 BGC in *O. ophidiicola* encodes a secondary metabolite known to inhibit squalene synthase ^56^ and may be a potential mechanism through which the fungus can modify its environment to promote colonization. Cholesterol had no effect on *O. ophidiicola,* perhaps due to it being the most abundant sterol in snake skin ^57^. The lipid composition of snake skin is both complex and variable and the effect of specific lipids range from no suppression to complete inhibition of *O. ophidiicola,* suggesting several lipids impart antifungal capabilities and/or *O. ophidiicola* is largely incapable of metabolizing these carbon sources. Based on these results, we expect lipid compounds widespread among snakes, and chosen for *in vitro* assays in this study, may underlie a potential antifungal defense system. Future work exploring species-specific differences in lipid chemistry may be a fruitful avenue in understanding SFD susceptibility ^58,59^.

Since the introduction of the chytrid pathogen of amphibians and white nose pathogen of bats, researchers have sought to characterize the skin microbiome response to fungal pathogenicity at the community-level in wild animals. A longitudinal laboratory inoculation study and broad scale survey of wild snakes provide strong evidence for pathogen induced dysbiosis by *O. ophidiicola* to the skin microbiome ^41,42^. Utilizing contrastive deep neural network models, we explored relationships between two different data domains and aligned the microbiome with the skin lipid profile of healthy and dysbiotic snakes. Our analytical approach extends beyond traditional analyses ^60^ and allows for targeted follow-up experiments exploring specific bacteria and lipids underlying BFIs and skin chemistry.

Through the integration of multifaceted *in vitro* assays and comparative genomic analyses, we were able to identify how BFIs influence fungal growth for four bacterial species: *Acinetobacter guillouiae, Chryseobacterium* sp.*, Flavobacterium odoratimimum,* and *Mammaliicoccus sciuri*. When *A. guillouiae* and *Chryseobacterium* sp. interacted directly with *O. ophidiicola*, they exhibited enhanced growth but only *Chryseobacterium* sp. inhibited fungal growth. Interestingly, the same isolate of *Chryseobacterium* sp. previously exhibited no fungal inhibitory activity when grown on tryptic soy agar (TSA) ^60^, suggesting that assays mimicking snake skin chemistry (keratin in this study) induce BGCs coding for secondary metabolites, such as flexirubin and siderophore production, both of which have known antifungal activity ^61^. Similarly, both *F. odoratimimum* and *M. sciuri* suppressed fungal growth, with the former having the BGC encoding flexirubin and the latter a siderophore BGC. *Mammaliicoccus sciuri* grew better in the absence of *O. ophidiicola*, and *F. odoratimimum* was unaffected by the fungus. Culturable bacteria from snake skin may be particularly enriched for siderophore biosynthesis and secondary metabolite production that dictate BFIs, a pattern that has been identified in other systems ^62,63^.

The bacterial metagenomes from free-ranging snakes largely recapitulated DBFI experiments. *Chryseobacterium* sp.*, F. odoratimimum,* and *M. sciuri* have growth-suppressive effects *in-vitro,* and were associated with disease negative snakes on the landscape, suggesting an important role in a healthy snake skin microbiome. Biosynthetic gene clusters for flexirubin and fulvivirgamide were identified in both the metagenomes and living bacterial isolates. Siderophores, including fulvivirgamide and desferrioxamine, were found in disease-negative snakes and may regulate metal ion availability ^64,65^, thus creating an environment less conducive for *O. ophidiicola* growth. The importance of iron availability in pathogenic fungi and their unique adaptations to exploit microbial pathways has been previously noted ^66,67^. *Ophidiomyces ophidiicola* contains a *Sit1* gene, which is a siderophore iron transporter that is associated with increased virulence in human-associated fungal pathogens. *Sit1* expression facilitates iron acquisition from neighboring bacteria and fungi ^67^, suggesting that *O. ophidiicola* may exploit bacterial siderophores to promote its own growth although this hypothesis requires further testing. The recapitulation of BFIs *in vitro,* and link to functional BGCs, scales from synthetic microbiomes to the ecosystem level, and suggests that important interactions can be characterized using simple reproducible assays to model dysbiosis.

Filamentous fungi grow by secreting exoenzymes to digest host resources and direct uptake of nutrients. Examining the effect of *O. ophidiicola* metabolism on bacterial growth highlighted how a fungal-altered environment, via keratin metabolism and metabolite production, can modulate the growth of skin bacteria. Dermatophyte pathogens closely related to *O. ophidiicola* metabolize keratin by reduction via sulphitolysis from excess sulfite released into the extracellular space and uptake short peptides and amino acids like cysteine ^68^. The reduced keratin then undergoes proteolysis through a variety of endo- and exopeptidases including subtilisin-like serine proteases and leucine aminopeptidases ^68^, both of which were identified in the *O. ophidiicola* genome. Primary metabolism of keratin by *O. ophidiicola* may result in cross-feeding interactions that initially support (5 day SM) *Chryseobacterium* sp. and *M. sciuri,* but later (9 day SM) suppress their growth due to reduction of small peptides and accumulation of secondary antimicrobial metabolites like chaetolivacine A ^69^. *Acinetobacter guillouiae* grew better on 9 day SM, perhaps due to its association with *O. ophidiicola* infected snakes, as represented in the metagenomics dataset, and known history as an opportunistic pathogen ^70^. Since we did not test the chemical composition of the SM directly, follow-up research is needed to identify which metabolites are produced by *O. ophidiicola,* and how they contribute to BFIs. Additionally, the use of a monoculture study design ignores the complex interactions that occur in bacterial communities that likely support diverse cross-feeding outcomes ^71^, especially in a complex carbon source like the skin ^72^.

To fully understand BFIs in the skin microbiome of snakes, both pathogen induced dysbiosis on bacterial assemblages and the effect of bacterial cross-feeding on *O. ophidiicola* should be considered. Through a series of cross-feeding experiments, we observed that fungal growth was reduced with the strongest effect on the bacterial SM only when bacterial cross-feeding interactions between *Chryseobacterium* and *S. maltophila* were tested. The antifungal compounds flexirubin and aryl polyene might explain this BFI. Members of the Onygenales, including *O. ophidiicola,* are known to exploit various byproducts of keratin proteolysis ^73^. The genome of *O. ophidiicola* has secreted subtilisin-like serine proteases which may allow for metabolism and growth on SM of *Chryseobacterium* and *S. maltophila* in monoculture. Scaling up experiments using enrichment cultures or more complex synthetic microbiomes may help reflect more complex species interactions ^74^ and allow for reproducible tests of dysbiosis.

The relationship between emerging fungal pathogens and the skin microbiome of afflicted hosts has received much attention, but studies largely ignore the complex BFIs and role of the skin environment in understanding fungal pathogen induced dysbiosis. Through a multifaceted approach combining landscape, laboratory, and *in vitro* experimental methods that incorporated multiple data domains, we were able to identify that snake skin lipids suppress the growth of *O. ophidiicola.* The inclusion of relevant host chemical composition into simple and reproducible experiments that explore host-microbiome-pathogen dynamics is an important step in understanding microbiome dysbiosis and disease ecology. The complex interplay in host-microbiome-pathogen systems highlight the need for similarly complex research projects that incorporate different data types to observe BFIs and explore underlying mechanisms of dysbiosis as we did in this study.

## Methods

### Overview of experimental approach

For all experiments, *O. ophidiicola* isolate 12-33400 was used (isolated from a *Thamnophis radix* by Allender et al. ^39^). A total of 26 bacterial isolates were obtained from free-ranging snakes (Supplementary Table 8; original source ^60^). The complete genomes of all 26 bacterial strains were sequenced, 11 of these isolates were included in the spent media experiments, and five of the 11 in the DBFI experiments. Eight isolates without full genomes were included in the spent media and DBFI experiments. The selection of bacterial isolates for each experiment was based on the deep learning models and attention scoring from Romer et al. ^41^ that predicted bacterial species associated with SFD state. All statistical analyses were conducted using R v4.3.2 ^75^, using two-tailed tests, and results are provided as β = effect size and *SE* = standard error. Additional methodology is provided in supplemental file 1.

### Snake skin lipid extraction and O. ophidiicola growth

Sheds from five snake species were collected after wild-caught individuals were brought into the lab or from fresh roadkill in Oregon, USA (Supplementary Table 1). Lipids were extracted from each snake skin (n = 6) after being submerged in a mixture of 1:1 dichloromethane:hexane solvent for 24 hours and then dried with nitrogen gas. The solution was then resuspended (15 mg lipids / 2 mL solvent) and divided into three technical replicates (n = 18) such that the lipid concentration in the medium for the described fungal assays was 1000 ppm (5 mg / 5 mL). To prepare the medium, 2 mL of the solvent with lipids was added to 15 mL of molten Sabouraud Dextrose Agar (SDA) using a lipid-free, autoclaved, glass pipet. Remaining solvent was evaporated under nitrogen gas, and lipids were evenly homogenized into the medium using a magnetic stir bar with 15 mL distributed evenly among three Petri dishes. A solvent control (solvent minus lipids) and negative control (only SDA) for each replicate were also prepared in triplicate. Plates were inoculated with a 6 mm plug of *O. ophidiicola*, sealed with Parafilm, and placed in an incubator at 24°C for 19 days. To quantify fungal growth, images of Petri dishes were captured and area of fungal growth (mm²) was determined in ImageJ ^76^. Any plates that exhibited bacterial contamination were removed from downstream analyses, resulting in a final sample size of 31. One-way ANOVAs were used to test significance, with or without Welch correction, depending on whether there was a violation of homogeneity of variance across treatments.

### Ophidiomyces ophidiicola growth on minimal media representative of snake skin chemistry

Keratin (Spectrum), cholesterol (VWR), oleic acid (TCI), and squalene (TCI) were prepared at five final concentrations (1, 10, 100, 1,000, 10,000 ppm), along with 200 mL of 5X M9 minimal salts, 15 g/L agar, and 2 mL of MgSO_4_ (1.0 M) and 0.1 mL of CaCl_2_ (1.0 M). Control plates included all the previous ingredients except with agarose as their only carbon source (herein called M9 control). After autoclaving and cooling, micronutrients were added, and each flask of media was continuously heated and stirred to ensure a homogenous concentration of lipids in each replicate (20 mL/plate). Plates were inoculated (n=5 replicates per treatment or control) with a 6mm plug of *O. ophidiicola* growing on M9 control plates (carbon negative), parafilmed, and incubated at 25°C for 22 days. The diameter of mycelial growth was measured across the plate in two perpendicular directions every three days for a total of 21 days. Additionally, we tested for an interactive effect of keratin and each lipid listed above using 10,000 ppm media.

Because *O. ophidiicola* exhibits linear growth *in vitro* ^39^ we utilized the diameter of fungal growth from the final date (day 21) as the response variable in all analyses. A linear model was used to determine the effect of carbon source and the interaction between carbon source and concentration, and significant pairwise differences were examined using a *post-hoc* Tukey test. Each carbon source was analyzed individually using a linear model to examine if fungal growth differed from M9 minimal media controls. The keratin plus lipid plates were compared to the 10,000 ppm keratin plates with a linear model (diameter ∼ medium) to determine how the lipid addition influenced fungal growth.

### Contrastive deep neural network to model skin microbiome and host chemistry

Juvenile Common Water Snakes (*Nerodia sipedon;* n = 22) were either inoculated with *O. ophidiicola* or a sham control and monitored over 80 days, or until an individual experienced mortality (see Romer et al. ^42^ for detailed methods). The skin microbiome of snakes was collected every seven days by swabbing a 15 cm portion of the midbody fifteen times with a rayon-tipped sterile applicator. DNA was extracted from swabs using the DNeasy PowerSoil kit (Qiagen) following the manufacturer’s protocol. A 250 bp region of the 16S V4 rRNA marker was amplified using the 515F and 806R primers and sequenced on a Illumina MiSeq. Bioinformatic analyses was performed in Mothur v1.43.0 ^77^ following the MiSeq protocol with minor modifications ^42^ and sequences were clustered into operational taxonomic units (OTUs) at 97% similarity. Upon the conclusion of the experiment, all snakes were euthanized and preserved in formalin.

To perform the skin lipid extraction, snake carcasses were rinsed of formalin with DI water for one minute before being placed into a 250 mL beaker filled with a 1:1 hexane:dichloromethane mix with the head and cloaca kept out of the beaker in order to avoid possible extraction of internal lipids. The beaker was then covered in aluminum foil to reduce evaporation. After a 24 hour period, the snake carcass was removed, solvent evaporated on a Rotavapor R3000, and the weight of the extracted lipids was measured.

Extracted lipids were prepared for gas chromatography mass spectrometry (GC/MS) by first converting acylglycerolipids to fatty acid methyl esters (FAMEs) ^78^ followed by converting any resulting hydroxy group-containing lipids to their trimethylsilyl (TMS) ether derivatives ^79^. GC/MS was carried out using the instrumentation and methodology detailed in ^78,80^, with the additions of two internal standards, sterane cholestane (mol. weight 372) and the linear hydrocarbon docosane (mol. weight. 310), both at 100 mg/L. Additionally, the final hold temperature of the GC (300 °C) was extended by 8 min. to a total run time of approximately 75 min. in order to accommodate potential late-eluting compounds (e.g., methyl ketones). Scanning was done from 50-650 *m/z* in positive-ion electron impact mode. GC/MS data acquired in the native .raw file format was converted to .mzML format using MSConvertGUI ^81^.

Contrastive deep learning methods were used to compare patterns in the skin microbiome to host lipid chemistry in the presence or absence of *O. ophidiicola* from the live animal study. Amplicon samples were processed using a set transformer ^82^ model composed of eight set attention blocks and eight attention heads. Given a set of DNA sequences, the model injects a learnable class token before passing everything through the transformers. The processed class token is conditioned on the input sequences and comes to represent the entire amplicon sample as a single latent vector embedding.

In order to process individual DNA sequence reads, we employ a modified version of the DNABERT model ^43^. Our DNABERT model consists of eight transformer blocks each with eight attention heads. We modified the architecture to employ relative position embeddings and removed the next sequence prediction task due to the short reads. The model was pre-trained for 100,000 steps on the SILVA v138.1 16S dataset ^44^ with a mask ratio of 15%, batch size of 256, and the Adam optimizer with a constant learning rate of 1e-4. Each batch consisted of sequences drawn uniformly at random, each randomly trimmed between 35 bp and 250 bp in length and encoded into 3-mer representations.

GC/MS samples were processed using a standard transformer model based on a vision transformer ^45^. The retention times were mapped directly to sinusoidal position encodings and total ion chromatography (TiC) signals were normalized to values ranging between 0 – 1. The encoded retention times and corresponding TiC signals were broken into non-overlapping patches of 20 points and linearly projected up to 256 dimensional embedding. A learnable class token was prepended to the patches and passed through eight transformer blocks each with eight attention heads. The class token after having passed through the transformers comes to represent the entire GC/MS sample as a single latent vector embedding.

Amplicon and GC/MS models were trained in tandem using contrastive pre-training for 30,000 steps. Batches consisted of six paired amplicon and GC/MS samples. Each amplicon subsample consisted of 1,000 uniformly drawn reads randomly truncated to 35-250 bp, and each GC/MS sample was randomly truncated on either end to 80-100% of its length (56 – 70 minutes) and then normalized. Each domain was processed by its corresponding model to generate embeddings. Models were trained following the contrastive pre-training objective as defined in^46^. In order to align the latent spaces, we shared the weights of the final linear projections and added an additional mean-squared error loss between the paired embeddings.

### Direct fungal-bacterial interactions

Six bacterial isolates (Supplementary Table 6) were streaked onto 10,000 ppm keratin minimal medium and grown at 25 °C for 48 hrs before beginning the experiment. 12-well microplates of M9 (only carbon source is agarose) or 10,000 ppm keratin were prepared by adding 5 mL agar minimal medium to each well. The following treatments and controls were set up in a fully factorial design including: 1) *O. ophidiicola* with/without a bacterial isolate on a 10,000 ppm keratin carbon source, 2) *O. ophidiicola* with/without a bacterial isolate on M9, and 3) bacterial isolate (no *O. ophidiicola*) on 10,000 ppm keratin carbon source or on M9. All 12-well plates had a respective medium blank without microbes to control for cross-contamination. The experimental design allowed for simultaneous tests of BFIs in response to both *O. ophidiicola* and host keratin. To inoculate the plates, a 4 mm diameter agar plug was removed from the center of each well, replaced with a plug of actively growing *O. ophidiicola* (or sterile M9 plug in bacteria only treatments) and held in place by adding 10 µL of molten M9 agar. An overlay of 1000 µL broth of 10,000 ppm keratin or M9 (control) was added to the matching 12-well plate location. A ten-fold serial dilution of bacterial isolates in phosphate buffered saline (PBS) was carried out in a 96-well flat bottom plate (Costar), measured on a ClarioStar plate reader at optical density (OD) 620 nm, and the dilution reading 0.15 nm used to inoculate the growth experiment below ^47^. To begin the experiment, 8 µL of each sample (0.15 nm cell density) was inoculated into the broth overlay in triplicate. The plates were incubated at 25 °C for 24 hrs then passaged six times, once every 24 hrs, by aspirating ∼1000 µL broth with bacterial growth and replenishment with fresh broth (1000 µL) into each well with keratin or M9. The standardized volume of remaining bacterial cells inoculated the fresh broth during each passage event. At each passage event, bacterial and fungal growth were simultaneously quantified by measuring OD at 620 nm for bacteria and *O. ophidiicola* radius of growth using a BioRad ChemiDoc MP Imaging System.

Linear regression was used to model the effect of time, media type, and bacterial and *O. ophidiicola* growth. Pairwise comparisons of estimated marginal means of linear trends were used to separate trend lines into statistically distinct groups. An ANOVA was used to model the effect of media type, *O. ophidiicola* presence, and interaction on area under the curve for bacterial growth calculated from OD620 nm measured every 24 hrs for six days. Area under the curve was calculated using Growthcurver ^83^. Since *Chryseobacterium* sp. was the only taxon with a significant interaction term, pairwise comparisons of estimated marginal means were used to determine differences within media types.

### Fungal-bacterial interactions along a nutrient/metabolite gradient

*Ophidiomyces ophidiicola* was grown on 1X M9 plates for 14 days at 25 °C. Ten 6 mm plugs of *O. ophidiicola* mycelia were added to three 2.8 L shaker flasks containing 1 L of 10,000 ppm keratin with 1X M9 salts broth minimal media and micronutrients (2 mL of MgSO4 [1.0 M]; 0.1 mL of CaCl2 [1.0 M]). The flasks were incubated at 25 °C while shaking at 120 rpm. To create a gradient of resource or metabolite usage by *O. ophidiicola*, a single shaker flask from each carbon source at days 5, 9, and 13 was filtered through autoclaved cheesecloth to remove the mycelial mass and collect the spent media (SM). Secondary vacuum filtration through a 0.2 µM pore size was used to remove the remaining fungal cells but retain the carbon source and metabolites. Frozen glycerol stocks of 18 strains of bacteria isolated from snakes (Supplementary Table 7) were thawed and passaged 3X times on 500 µL of a 10,000 ppm keratin medium to revive the culture then plated out on 10,000 ppm keratin plates. To begin the growth rate experiments, a serial dilution was conducted by using a sterile needle to pick up several colonies from each plate and inoculate them into 275 µL of sterile PBS. A ten-fold serial dilution was carried out, OD620 nm measured, and the dilution reading 0.15 used to inoculate the growth experiment below ^47^. Volumes of 2 µL of each sample (0.15 cell density) were inoculated in triplicate into 250 µL of 10,000 ppm keratin minimal media (positive control) and 5, 9 or 13 day SM. All experiments were conducted in 96-well flat bottom plates (Costar) inside of a laminar flow hood and measured on a ClarioStar plate reader using an absorbance reading at OD620 nm. Plates were incubated at 25 °C and absorbance reading values were recorded each hour for 50 hrs to measure bacterial growth. The plate was shaken at 200 rpm for 30 secs before each measurement and the well was scanned using the spiral (5mm) average setting. Differences in bacterial growth were determined with an ANOVA using area under the curve as the response variable with media as the predictor variable.

*Chryseobacterium* sp. (isolate BR1.10) and *Stenotrophomonas maltophilia* (isolate TR087-6.4) were grown on 10,000 ppm keratin plates and a single colony was selected, serially diluted using the protocol above, then inoculated (2 µL at 0.15 - 600 nm) into 100 mL of 10,000 ppm keratin or M9 control broth and incubated at 25 °C while shaking (100 rpm in 250 mL shaker flasks). After 24 hrs, each of the six broth cultures (species + keratin, keratin control, species + M9, M9 control) were passed through a 0.22 uM filter to remove the bacterial cells and collect the SM. The SM was partitioned into two aliquots including 60 mL for *O. ophidiicola* growth assays and 40 mL for bacterial cross-feeding experiments. *Chryseobacterium* sp. or *S. maltophilia* were serially diluted and inoculated as above into 40 mL of the opposite strain’s SM in a 75 mL flask then incubated at 25 °C while shaking (100 rpm). After 24 hrs of growth the media were filter sterilized and stored at 4 °C for four days. The liquid SM was added in an equal volume to a molten (∼60 °C) 2X agar medium, under constant mixing, then 10 mL dispensed into four replicate 60 mm Petri plates per treatment or control to test for 1) the effect of independent bacterial strains on *O. ophidiicola* growth, 2) additive effects by combining the SM of each bacterial strain, and 3) cross-feeding of the SM from both bacterial strains and the impact on *O. ophidiicola* growth. The diameter of fungal growth on bacterial SM was measured every two days over 14 days. For the analysis, fungal growth on day 14 was the response variable with medium and bacterial treatment as predictor variables and plate replicate as a random effect. Single bacterium and mixed bacteria treatments were separated in the analysis.

### Genomic predictions of interactions between skin bacteria and O. ophidiicola

To predict putative gene clusters associated with the production of secondary metabolites that may play a role in host-microbiome-pathogen interactions, the fasta file of the reference genome of *O. ophidiicola* (NCBI RefSeq accession number GCF_022830035.1) was used in conjunction with the annotation .gff file to identify biosynthetic gene clusters (BGCs) with antiSMASH version 7.1 ^84^.

To understand putative skin microbiome function of snakes on the landscape, bacterial metagenomes were sequenced from 120 wild Common Watersnakes. To overcome issues with low yield of DNA from skin swabs, we pooled n=10 DNA isolations based on disease status for snakes at a particular site in Tennessee, USA. In total, we prepared n=4 pooled libraries of *O. ophidiicola* positive snakes and n=8 negative libraries using the Nextera DNA Flex kit. The libraries were sequenced on an Illumina NextSeq 500 (2 x 75 bp paired end reads).

Raw sequencing reads underwent quality control using Trimmomatic (v0.39) ^85^ to remove adapter sequences, low-quality bases, and short reads. To recover metagenome-assembled genomes (MAGs), three binning algorithms were utilized: MaxBin2 ^86^, MetaBAT2 ^87^, and CONCOCT ^88^. The outputs from the three algorithms were consolidated using DASTool ^89^ to generate final high-quality bins for each sample. Completeness and contamination of the resulting bins was assessed using CheckM (v1.1.2) ^90^. Bins were classified as high quality (>90% completeness) or draft quality (70–90% completeness) only if contamination levels were below 5% dRep (v2.6.0) ^91^. Redundant bins were removed to ensure the retention of the most representative genomes for taxonomic classification and downstream analyses.

Dereplicated MAGs were taxonomically classified with GTDB-Tk (v2.3.2) ^92^, following the Genome Taxonomy Database (GTDB) release 214. Only genomes with high completeness (>90%) and low contamination (<5%) were included in subsequent phylogenetic and functional analyses. Anvi’o^93^ Q2Q3 data were used to estimate the abundance of each bin across the metagenomic samples. Abundance data were normalized by the total number of base pairs in the paired-end reads using a per-gigabase scaling approach to ensure abundance values accurately reflected microbial community composition across the samples.

Genomes of 26 bacterial strains (Supplementary Table 8) from host species were isolated from snakes. High-molecular-weight genomic DNA was extracted using the MasterPure Yeast DNA Purification Kit (Biosearch Technologies Genomic Analysis by LGC MPY80200) from the sucrose-washed bacterial pellets according to manufacturer’s specifications and sent to SeqCoast Genomics for short read whole genome sequencing with Illumina or long read sequencing with Oxford Nanopore Technologies.

Raw Illumina sequencing reads underwent initial quality control, filtering, and adapter trimming using fastp v0.23.2 ^94^. After preprocessing, the high-quality paired-end reads were assembled de novo into contiguous sequences (contigs) using SPAdes v3.15.5 ^95^. The resulting assemblies were analyzed for BGCs using antiSMASH v6.1.1 ^96^, a tool for the detection and annotation of secondary metabolite gene clusters in microbial genomes. Each identified BGC was then manually inspected to ensure accurate characterization and to enable detailed comparisons against a database of previously described BGCs available in the Minimum Information about a Biosynthetic Gene Cluster (MIBiG) repository, version 4.0rc1 ^97^. This step allowed for the identification of known or novel biosynthetic pathways, facilitating further exploration of their potential for molecules involved in SFD microbiome interactions.

## Acknowledgements

This work was supported by National Science Foundation grants EF-2125065 and CAREER 2236580 to D. Walker, the Molecular Biosciences Program at MTSU, TWRA State Wildlife Grant #58209 to D. Walker, and National Science Foundation grants REU 1757493 and DEB 1933925 to J. Phillips. C. Van Moorleghem was funded through the Fulbright Visiting Scholar Program. M. G. Chevrette and M. S. Chue Donahey were funded through the Department of Microbiology and Cell Science at the University of Florida and USDA awards 58 6066 4 047 and 6066 21310 005 052 S. All research was conducted under IACUC19-3012, IACUC 19-3001, TWRA1547, TWRA1907, TDEC 2016-026. The authors would like to thank Alexander Romer, Kylie Moe and Reed Alexander for their technical assistance and discussions around this project.

## Competing interests

The authors declare no competing financial interests.

## Data availability statement

Illumina amplicon reads analysed in this study have been deposited in GenBank under BioProject accession number PRJNA1114724 (https://www.ncbi.nlm.nih.gov/bioproject/1114724) and PRJNA1114659 (https://www.ncbi.nlm.nih.gov/bioproject/1114659). Datasets generated during the current study are available at https://github.com/DLii-Research/sfd-skin-chemistry

## Code Availability

R code and contrastive deep learning models are available at https://github.com/DLii-Research/sfd-skin-chemistry.

## Author Contributions

Conceptualization: D.M.W., Methodology: J.L.P., D.L., J.H.N., D.M.W., Software: J.L.P., D.L., T.S., Formal Analysis: K.M.M., J.W.D., L.V.G., C.V.M., J.L.P., D.L., T.S., J.H.N., J.L., C.S., O.B., M.S.C.D., M.G.C., D.M.W., Investigation: K.M.M., J.W.D., L.V.G., C.V.M., J.H.N., J.L., J.M.M., C.S., O.B., M.S.C.D., M.G.C., D.M.W., Resources: J.L.P., J. L., J.M.M., M.G.C., D.M.W., Data Curation: K.M.M., J.W.D., D.M.W., Writing – Original Draft: K.M.M., J.W.D., L.V.G., J.L.P., D.L., M.R.P., D.M.W., Writing - Review & Editing: K.M.M., J.W.D., C.V.M., M.R.P., M.G.C., D.M.W., Visualization: K.M.M. J.W.D., J.L.P., D.L., D.M.W., Supervision: R.M., J.W.S., D.M.W., Project administration: D.M.W., Funding acquisition: J.L.P., D.M.W.

## Supplementary Info

**Supplementary File 1:** Detailed description of all materials and methods.

**Supplementary Table 1:** Biographics of snake skins utilized in Fig. 1.

**Supplementary Table 2:** Output of a linear regression model examining the diameter of *Ophidiomyces ophidiicola* growth (mm) after 21 days on four types of minimal media (keratin, cholesterol, oleic acid, and squalene) and five concentrations (10,000, 1,000, 100, 10, and 1 ppm). For the model, the keratin medium and 10,000 ppm concentration were set as reference levels.

**Supplementary Table 3:** Pairwise contrasts using Tukey’s HSD accounting for significant interactions in the linear regression model from Supplementary Table 3. Estimates shown are *Ophidiomyces ophidiicola* diameter (mm).

**Supplementary Table 4:** Linear regression models examining the diameter of *Ophidiomyces ophidiicola* growth (mm) within each of the four minimal media treatments relative to M9 minimal media controls. Variables shown represent the medium and its concentration (i.e., Keratin 10,000 represents Keratin at 10,000 ppm).

**Supplementary Table 5:** Linear regression models examining the diameter of *Ophidiomyces ophidiicola* growth (mm) across mixed media plates with 10,000 ppm keratin as the reference level.

**Supplementary Table 6:** Growth of bacterial species (isolate ID given in parentheses), in optical density, in the presence and absence of *O. ophidiicola* (*Oo*) grown in co-culture using an ANOVA. * = growth of *Oo* in the presence of each bacterial isolate (in unit) on keratin media across time using a linear fixed effects model.

**Supplementary Table 7:** Pairwise contrasts using Tukey’s HSD of bacterial growth across the different media types in the fungal spent media experiment.

**Supplementary Table 8:** Putative biosynthetic gene clusters from 26 bacterial isolates obtained from free-ranging snakes. Inhibits *Oo* status represents *in vitro* assay results from Hill et al. ^60^.

**Supplementary Table 9:** Diversity of putative biosynthetic gene clusters identified in the *Ophidiomyces ophidiicola* genome.

---

**Supplementary Fig. 1.**
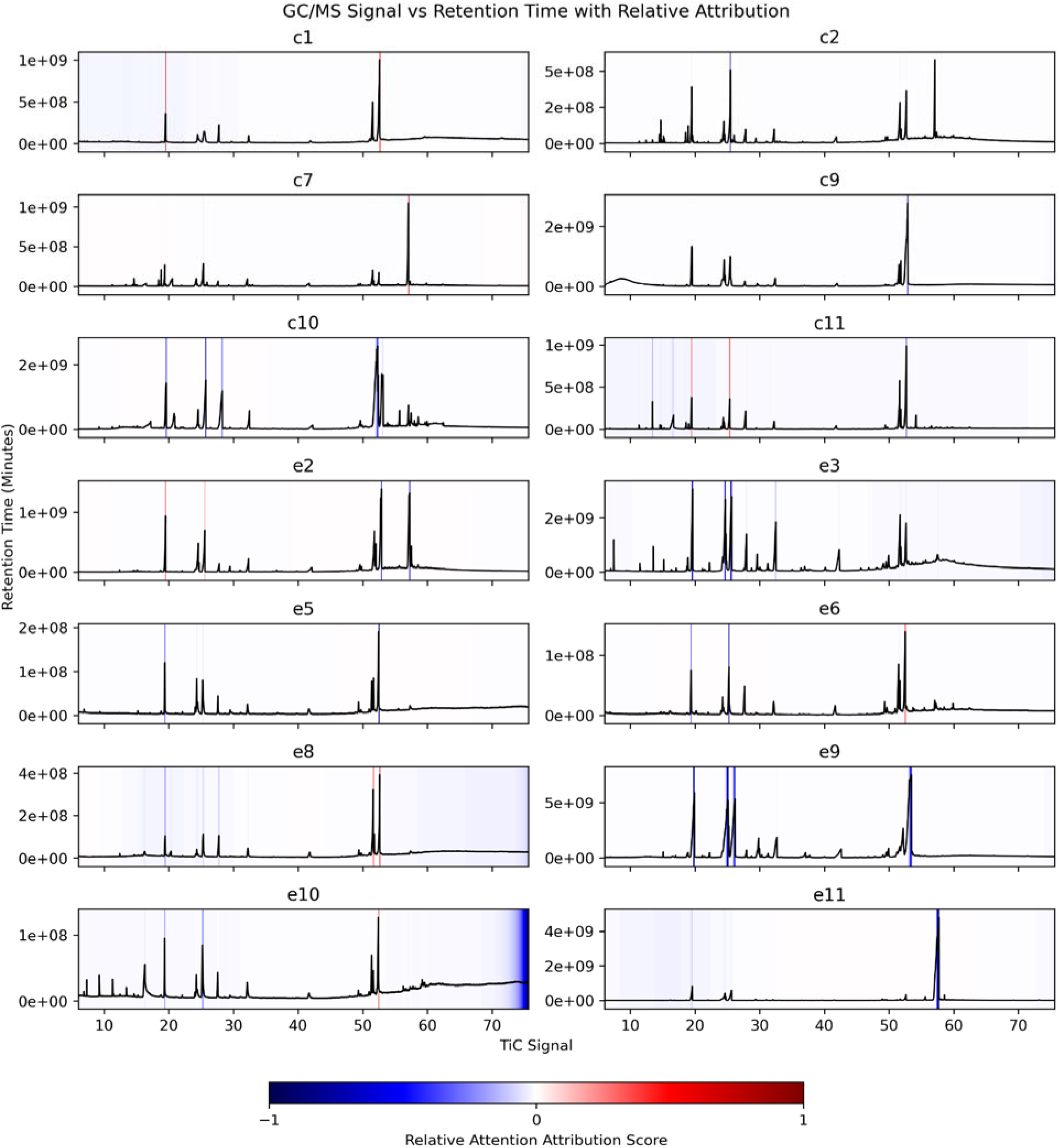
Deep neural network attention scores of snake skin GC/MS profiles. The GC/MS results from a live animal study of common water snakes (*Nerodia sipedon*) experimentally infected with *Ophidiomyces ophidiicola* (plots denoted by eX) and control individuals (plots denoted by cX). The total ion chromatography (TiC) signals are color coded with normalized relative attribution scores. For the control panels, red indicates peaks/regions that drive negative infection status prediction while blue regions hinder prediction. For the experimental panels, red indicates peaks/regions that drive positive infection status prediction while blue regions hinder prediction.

**Supplementary Fig. 2.**
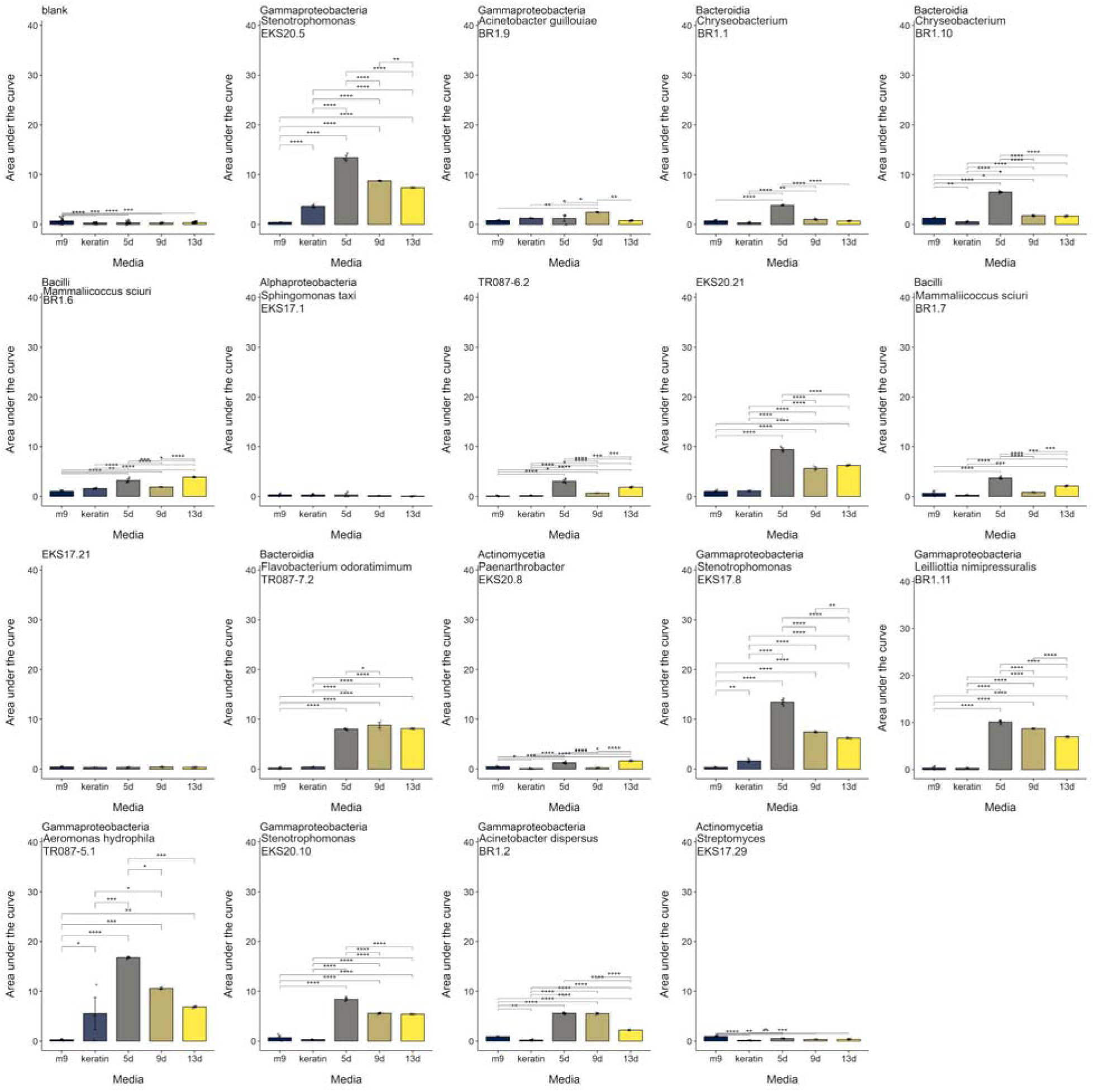
The growth of bacteria on *Ophidiomyces ophidiicola* metabolized keratin. The effect of *O. ophidiicola* SM at 0 (“keratin”), 5, 9, or 13 days of fungal metabolism of a 10,000 ppm keratin medium across all bacterial isolates. Statistically significant pairwise differences via a Tukey HSD are indicated by: p < 0.05, *; p < 0.01, **; p < 0.001, ***; and p < 0.0001, ****. Error bars denote ± 1 standard error.

**Supplementary Fig. 3.**
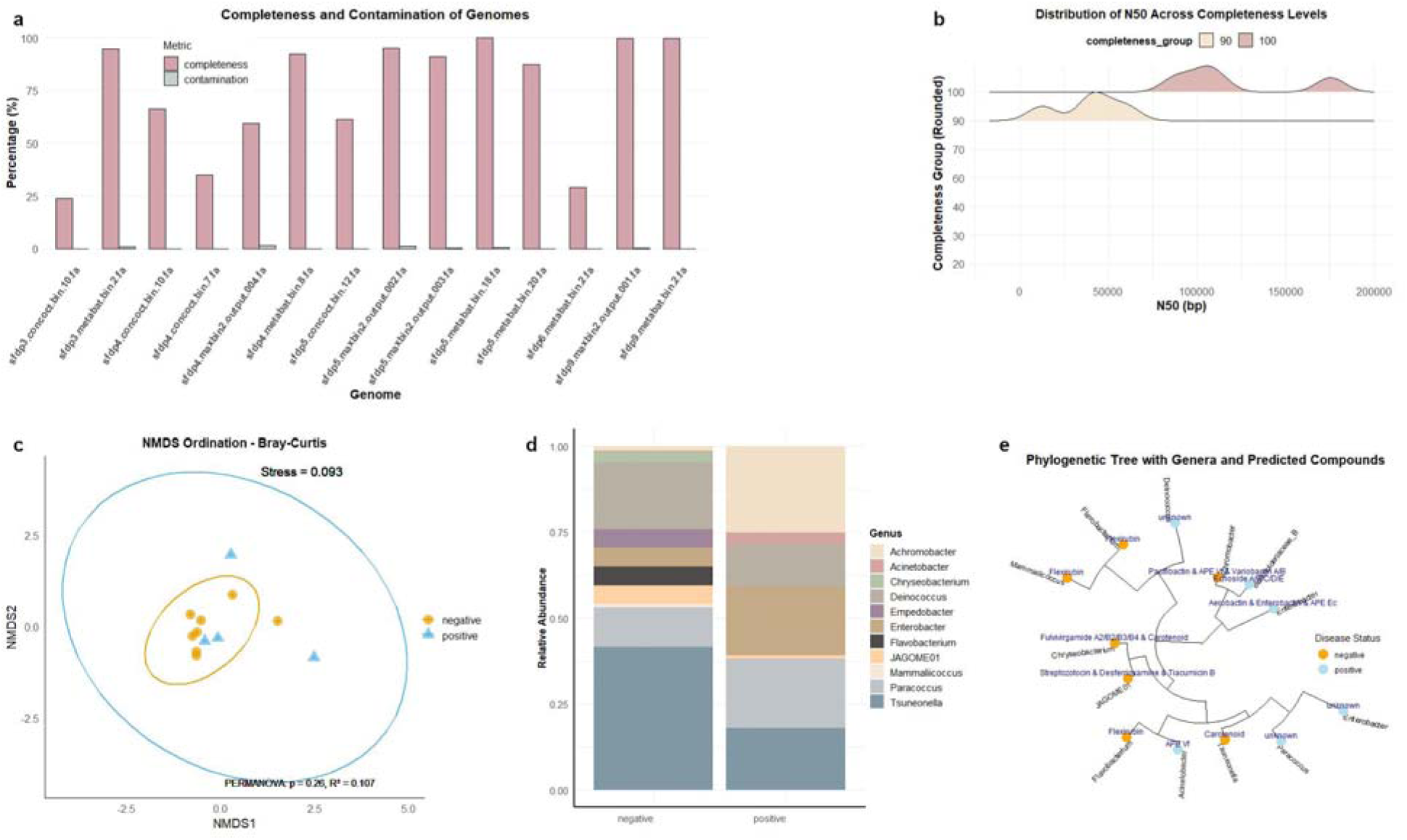
Metagenome-assembled genomes (MAGs) from wild snake skin microbiomes. a) Completeness and contamination assessment of MAGs. The completeness (pink bars) and contamination (brown bars) were evaluated for each bacterial MAG, with most showing high completeness and low contamination. b) Density plot of N50 values categorized by genome completeness. Genomes with higher completeness percentages (90% and 100%) tend to exhibit higher N50 values. c) NMDS ordination plot based on Bray-Curtis dissimilarity showing MAGs differences between disease status groups (negative vs. positive). The clustering indicates differences, with stress value and PERMANOVA results suggesting potential group separations. d) Stacked bar plot showing the relative abundance of bacterial genera in negative and positive disease groups. e) Phylogenetic tree of bacterial genera with predicted metabolic compounds. Genera associated with disease status (negative vs. positive) are indicated.

## Notes

https://doi.org/10.6084/m9.figshare.30199834.v1

https://www.ncbi.nlm.nih.gov/bioproject/1114724

https://www.ncbi.nlm.nih.gov/bioproject/1114659

https://doi.org/doi:10.25345/C59K4661B

### Competing Interest Statement

The authors have declared no competing interest.

