## Supplementary File 1 for "Skin lipid chemistry influences host-microbiome-pathogen interactions in snake fungal disease (ophidiomycosis)"

**Methods**

*Overview of experimental approach*

For all experiments, *O. ophidiicola* isolate 12-33400 was used (isolated from a *Thamnophis radix* by Allender et al. ^1^). A total of 26 bacterial isolates were obtained from free-ranging snakes (Supplementary Table 8; original source ^2^). The complete genomes of all 26 bacterial strains were sequenced, 11 of these isolates were included in the spent media experiments, and five of the 11 in the DBFI experiments. Eight isolates without full genomes were included in the spent media and DBFI experiments. The selection of bacterial isolates for each experiment was based on the deep learning models and attention scoring from Romer et al. ^3^ that predicted bacterial species associated with SFD state. All statistical analyses were conducted using R v4.3.2 ^4^ and results are provided as *β* = effect size and *SE* = standard error.

*Snake skin lipid extraction and O. ophidiicola growth*

From 26 March to 15 August 2021, sheds from five snake species (*Pituophis catenifer*, *Diadophis punctatus*, *Thamnophis sirtalis*, *Crotalus oreganus*, and *Charina bottae*) were collected after wild-caught individuals were brought into the lab or from fresh roadkill in Oregon, USA (Supplementary Table 1). Upon collection, sheds were stored at 4°C until lipid extraction. Lipids were extracted from each snake skins (n = 6) after being submerged in a mixture of 1:1 dichloromethane:hexane solvent for 24 hours and then dried with nitrogen gas. The solution was then resuspended (15 mg lipids / 2 mL solvent) and divided into three technical replicates (n = 1817) such that the lipid concentration in the medium for the described fungal assays was 1000 ppm (5 mg / 5 mL). To prepare the medium, 2 mL of the solvent with lipids was added to 15 mL of molten Sabouraud Dextrose Agar (SDA) using a lipid-free, autoclaved, glass 5 mL pipet. Remaining solvent was evaporated under nitrogen gas, and lipids were evenly homogenized into the medium using a magnetic stir bar. The lipid-containing medium (5 mL) was poured into three Petri dishes. A solvent control (solvent minus lipids) and negative control (only SDA) for each replicate were also prepared in triplicate. Plates were inoculated with a 6 mm plug of *O. ophidiicola*, sealed with Parafilm, and placed in an incubator at 24°C for 19 days. To quantify fungal growth, images of Petri dishes were captured and area of fungal growth (mm²) was determined in ImageJ ^5^. Any plates that exhibited bacterial contamination were removed from downstream analyses, resulting in a final sample size of 31. One-way ANOVAs were used to test significance, with or without Welch correction, depending on whether there was a violation of homogeneity of variance across treatments.

*Ophidiomyces ophidiicola growth on minimal media representative of snake skin chemistry*

Keratin, cholesterol, oleic acid, and squalene are some of the most abundant compounds present in snake skin^6^ and were thus used to test for *O. ophidiicola* growth facilitation or inhibition in carbon minimal media. Keratin (Spectrum), cholesterol (VWR), oleic acid (TCI), and squalene (TCI) were prepared at five final concentrations (1, 10, 100, 1,000, 10,000 ppm), along with 200 mL of 5X M9 minimal salts, 15 g/L agar, and 2 mL of MgSO_4_ (1.0 M) and 0.1 mL of CaCl_2_ (1.0 M). Control plates included all of the previous ingredients except with agarose as their only carbon source (herein called M9 control). After autoclaving and cooling, micronutrients were added, and each flask of media was continuously heated and stirred to ensure a homogenous concentration of lipids in each replicate (20 mL/plate). Plates were inoculated (n=5 replicates per treatment or control) with a 6mm plug of *O. ophidiicola* growing on M9 control plates (carbon negative)*,* parafilmed, and incubated at 25°C for 22 days under a 12:12 light-dark cycle. The diameter of mycelial growth was measured across the plate in two perpendicular directions every three days for a total of 21 days. Additionally, we tested for an interactive effect of keratin and each lipid listed above using 10,000 ppm media.

Because *O. ophidiicola* exhibits linear growth *in vitro* ^1^ we utilized the diameter of fungal growth from the final date (day 21) as the response variable in all analyses. A linear model (diameter ~ medium + medium x concentration) was used to determine the effect of carbon source and interaction between carbon source and concentration and assessed via a two-tailed ANOVA. Significant pairwise differences were examined *post-hoc* using ‘*emmeans::contrast’* to account for the interaction term ^7^. Each carbon source was analyzed individually using a linear model to examine if fungal growth on these plates differed from M9 minimal media controls. The keratin plus lipid plates were compared to the 10,000 ppm keratin plates with a linear model (diameter ~ medium) to determine how the lipid addition influenced fungal growth.

*Direct fungal-bacterial interactions*

Direct bacterial-fungal interaction experimental setup was a three-step process including i) growth of *O. ophidiicola*, ii) preparation of bacterial isolates, and iii) plate setup and passaging events. We grew *O. ophidiicola* on M9 minimal medium plates for 12 days at 25 °C prior to experimental plate inoculation. Six bacterial isolates (Supplementary Table 6) were streaked onto 10,000 ppm keratin minimal medium and grown at 25 °C for 48 hrs before beginning the experiment. 12-well microplates of M9 (only carbon source is agarose) or 10,000 ppm keratin were prepared by adding 5 mL agar minimal medium to each well. The following treatments and controls were set up in a fully factorial design including: 1) *O. ophidiicola* with/without a bacterial isolate on a 10,000 ppm keratin carbon source, 2) *O. ophidiicola* with/without a bacterial isolate on M9 (minus host carbon source), and 3) bacterial isolate (no *O. ophidiicola*) on 10,000 ppm keratin carbon source or on M9 (minus host carbon source). All 12-well plates had a respective medium blank without microbes to control for plate cross-contamination. The experimental design allowed for simultaneous tests of bacterial-fungal interactions (BFI) in response to both *O. ophidiicola* and host keratin. To inoculate the plates, a 4 mm diameter agar plug was removed from the center of each well, replaced with a plug of actively growing *O. ophidiicola* (or sterile M9 plug in bacteria only treatments) and held in place by adding 10 µL of molten M9 agar. An overlay of 1000 µL broth of 10,000 ppm keratin or M9 (control) was added to the matching 12-well plate location. A ten-fold serial dilution of bacterial isolates in phosphate buffered saline (PBS) was carried out in a 96-well flat bottom plate (Costar), measured on a ClarioStar plate reader at optical density 620 nm, and the dilution reading 0.15 nm used to inoculate the growth experiment below ^19^. To begin the experiment, 8 µL of each sample (0.15 nm cell density) was inoculated into the broth overlay in triplicate. The plates were incubated at 25 °C for 24 hrs then passaged six times, once every 24 hrs, by aspirating ~1000 µL broth with bacterial growth and replenishment with fresh broth (1000 µL) into each well with keratin or M9. The standardized volume of remaining bacterial cells inoculated the fresh broth during each passage event. At each passage event, bacterial and fungal growth were simultaneously quantified by measuring optical density (OD) at 620 nm for bacteria and *O. ophidiicola* radius of growth using a BioRad ChemiDoc MP Imaging System to quantify the radius of fungal extension.

Linear regression was used to model the effect of time, media type, and bacterial and *O. ophidiicola* growth. Pairwise comparisons of estimated marginal means of linear trends were used to separate trend lines into statistically distinct groups. A two-tailed ANOVA was used to model the effect of media type, *O. ophidiicola* presence, and interaction on area under the curve for bacterial growth calculated from OD620 nm measured every 24 hrs for six days. Area under the curve was calculated using Growthcurver ^20^. Since *Chryseobacterium* sp. was the only taxon with a significant interaction term, pairwise comparisons of estimated marginal means were used to determine differences within media types.

*Fungal-bacterial interactions along a nutrient/metabolite gradient*

To explore the indirect effects of a fungal modified environment on bacterial growth, keratin media metabolized by *O. ophidiicola* was used*,* as bacterial communities are known to respond directly to fungal competition for a nutrient source or fungal-derived exometabolites ^21^*. Ophidiomyces ophidiicola* was grown on 1X M9 plates for 14 days at 25 °C with no carbon source aside from agarose. Ten 6 mm plugs of *O. ophidiicola* mycelia were added to three 2.8 L shaker flasks containing 1 L of 10,000 ppm keratin with 1X M9 salts broth minimal media contam(single carbon source) and micronutrients (2 mL of MgSO4 [1.0 M]; 0.1 mL of CaCl2 [1.0 M]). The flasks were incubated at 25 °C while shaking at 120 rpm. To create a gradient of resource or metabolite usage by *O. ophidiicola*, a single shaker flask from each carbon source at days 5, 9, and 13 was filtered through autoclaved cheesecloth to remove the mycelial mass and collect the spent media (SM). Secondary vacuum filtration through a 0.2 µM pore size was used to remove the remaining fungal cells, but retain the carbon source and metabolites. The *O. ophidiicola* SM was used in subsequent experiments described below. Frozen glycerol stocks of 18 strains of bacteria isolated from snakes (Supplementary Table 8) were thawed and passaged 3X times on 500 µL of a 10,000 ppm keratin medium to revive the culture then plated out on 10,000 ppm keratin plates. To begin the growth rate experiments, a serial dilution was conducted by using a sterile needle to pick up several colonies from each plate and inoculate them into 275 µL of sterile PBS. As in the above DBFI experiment, a ten-fold serial dilution was carried out, OD620 nm measured, and the dilution reading 0.15 used to inoculate the growth experiment below ^19^. Volumes of 2 µL of each sample (0.15 cell density) were inoculated in triplicate into 250 µL of 10,000 ppm keratin minimal media (positive control) and 5, 9 or 13 day SM. As additional controls, each strain was inoculated in triplicate into 1X M9 to confirm a growth response solely to nutrients or metabolites, but not M9 minimal salts. On each plate, control blanks included sterile 1X M9 and 10,000 ppm keratin wells in triplicate to account for cross-well contamination. All experiments were conducted in 96-well flat bottom plates (Costar) inside of a laminar flow hood and measured on a ClarioStar plate reader using an absorbance reading at OD620 nm. Plates were incubated at 25 °C and absorbance reading values were recorded each hour for 50 hrs to measure bacterial growth. The plate was shaken at 200 rpm for 30 secs before each measurement and the well was scanned using the spiral (5mm) average setting. Differences in bacterial growth were determined with a two-tailed ANOVA using area under the curve as the response variable with media as the predictor variable.

Raw sequencing reads underwent quality control using Trimmomatic (v0.39) ^23^ to remove adapter sequences, low-quality bases, and short reads. Reads with Phred scores below 30 were trimmed using a sliding window approach (50:30), and reads shorter than 50 bp were discarded. Post-trimming, read quality was assessed using FastQC ^24^ to ensure the removal of low-quality regions and adapter contamination (Supplementary Fig. 3). High-quality paired-end and unpaired reads were assembled using SPAdes (v3.11.1) ^25^, applying the following k-mer values: 21, 33, 55, 77, 99, and 127. The options –meta, –sc, and –careful were employed to optimize the assembly for metagenomic datasets. Contigs shorter than 5,000 bp were excluded from the assembly to retain longer and more informative sequences. Anvi’o (v8) ^26^ was then used to reformat the contigs, ensuring the dataset contained only high-confidence contigs (< 90% contamination) for downstream analysis. Trimmed reads were mapped back to the assembly using Bowtie2 (v2.3.4.3) ^27^, and SAM files were converted to sorted BAM files using Samtools (v1.9) ^28^. Anvi’o was employed to generate contig databases and profile the samples, enabling the computation of coverage and detection of single-copy core genes across the metagenomes. To recover metagenome-assembled genomes (MAGs), three binning algorithms were utilized: MaxBin2 ^29^, MetaBAT2 ^30^, and CONCOCT ^31^. Each algorithm independently assigned contigs to bins: MaxBin2 utilized abundance profiles, while MetaBAT2 and CONCOCT incorporated both abundance and tetranucleotide frequency information. The outputs from the three algorithms were consolidated using DASTool ^32^ to generate final high-quality bins for each sample. Completeness and contamination of the resulting bins was assessed using CheckM (v1.1.2) ^33^. Bins were classified as high quality (>90% completeness) or draft quality (70–90% completeness) only if contamination levels were below 5% dRep (v2.6.0) ^34^. Redundant bins were removed to ensure the retention of the most representative genomes for taxonomic classification and downstream analyses.

Dereplicated MAGs were taxonomically classified with GTDB-Tk (v2.3.2) ^35^, following the Genome Taxonomy Database (GTDB) release 214. Only genomes with high completeness (>90%) and low contamination (<5%) were included in subsequent phylogenetic and functional analyses. A multi-locus phylogenomic tree of the recovered MAGs was constructed using FastTreeMP (v2.1.11) ^36^. Anvi’o Q2Q3 data were used to estimate the abundance of each bin across the metagenomic samples. Abundance data were normalized by the total number of base pairs in the paired-end reads using a per-gigabase scaling approach to ensure abundance values accurately reflected microbial community composition across the samples. These profiles were utilized in downstream statistical analyses to assess microbial community structure and functional capacity.

Genomes of 26 bacterial strains (Supplementary Table 8) from host species were isolated from snakes. Single colonies were picked from each plate and streaked onto tryptic soy agar to ensure isolation. The plates were incubated at 28 °C for 18 hours. For each strain, 2 mL of tryptic soy broth was inoculated with an isolated colony. The cultures in suspension were incubated at 28 °C with shaking at 250 rpm until turbid then centrifuged at 3,220 rcf for 5 minutes. Without disturbing the bacterial pellets, the supernatant in each tube was decanted. To remove media and periplasmic proteins, the bacterial pellets were washed in 300mM sucrose that had been sterilized with a 0.2-micron polyethersulfone (PES) filter. The bacterial pellets were resuspended in 1 mL of 300 mM sucrose and suspensions were centrifuged at 3,220 rcf for 5 minutes. Without disturbing the bacterial pellets, the supernatant in each tube was decanted. The process was repeated for a total of two washes. Then the bacterial pellets were frozen at -20 °C for a minimum of 20 minutes to aid cellular lysing and to provide a storage temperature with minimal enzymatic activity. High-molecular-weight genomic DNA was extracted using the MasterPure Yeast DNA Purification Kit (Biosearch Technologies Genomic Analysis by LGC MPY80200) from the sucrose-washed bacterial pellets according to manufacturer's specifications (see Supplemental Methods for minor deviations) and sent to SeqCoast Genomics for short read whole genome sequencing with Illumina or long read sequencing with Oxford Nanopore Technologies.

Raw Illumina sequencing reads underwent initial quality control, filtering, and adapter trimming using fastp v0.23.2 ^37^. After preprocessing, the high-quality paired-end reads were assembled de novo into contiguous sequences (contigs) using SPAdes v3.15.5 ^38^. The resulting assemblies were analyzed for BGCs using antiSMASH v6.1.1 ^39^, a tool for the detection and annotation of secondary metabolite gene clusters in microbial genomes. Each identified BGC was then manually inspected to ensure accurate characterization and to enable detailed comparisons against a database of previously described BGCs available in the Minimum Information about a Biosynthetic Gene Cluster (MIBiG ) repository, version 4.0rc1 ^40^. This step allowed for the identification of known or novel biosynthetic pathways, facilitating further exploration of their potential for molecules involved in SFD microbiome interactions.
