## Supplementary Table 1 for "Skin lipid chemistry influences host-microbiome-pathogen interactions in snake fungal disease (ophidiomycosis)"

**Supplementary Table 1:** Biographics of snake skins utilized in Fig. 1.

| Extraction ID | Species | Subspecies | Material type | Date of collection |
| --- | --- | --- | --- | --- |
| CB4S0719 | *Charina bottae* |  | Shed | 7/19/2021 |
| CO3D0815 | *Crotalus oreganus* |  | Salvaged (roadkill) | 8/15/2021 |
| DP14S0601 | *Diadophis punctatus* |  | Shed | 6/1/2021 |
| PCC13D0630 | *Pituophis catenifer* | *deserticola* | Salvaged (roadkill) | 6/30/2021 |
| PCC14D0705 | *Pituophis catenifer* | *catenifer* | Salvaged (roadkill) | 7/5/2021 |
| TSF1S0701 | *Thamnophis sirtalis* | *fitchi* | Shed | 7/1/2021 |
