## Supplementary Table 2 for "Skin lipid chemistry influences host-microbiome-pathogen interactions in snake fungal disease (ophidiomycosis)"

**Supplementary Table 2:** Output of a linear regression model examining the diameter of *Ophidiomyces ophidiicola* growth (mm) after 21 days on four types of minimal media (keratin, cholesterol, oleic acid, and squalene) and five concentrations (10,000, 1,000, 100, 10, and 1 ppm). For the model, the keratin medium and 10,000 ppm concentration were set as reference levels.

| *Predictors* | *Estimates* | *CI* | *p* |
| --- | --- | --- | --- |
| (Intercept) | 62.05 | 60.23 – 63.87 | **<0.001** |
| MEDIUM [Cholesterol] | -19.2 | -21.78 – -16.62 | **<0.001** |
| MEDIUM [Oleic Acid] | -44.9 | -47.48 – -42.32 | **<0.001** |
| MEDIUM [Squalene] | -62.05 | -64.63 – -59.47 | **<0.001** |
| CONC [1,000] | -6.25 | -8.83 – -3.67 | **<0.001** |
| CONC [100] | -10.05 | -12.63 – -7.47 | **<0.001** |
| CONC [10] | -22.36 | -25.10 – -19.63 | **<0.001** |
| CONC [01] | -14.2 | -16.78 – -11.62 | **<0.001** |
| MEDIUM [Cholesterol] × CONC [1,000] | 4.25 | 0.60 – 7.90 | **0.023** |
| MEDIUM [Oleic Acid] × CONC [1,000] | 8.75 | 5.10 – 12.40 | **<0.001** |
| MEDIUM [Squalene] × CONC [1,000] | 6.25 | 2.49 – 10.01 | **0.001** |
| MEDIUM [Cholesterol] × CONC [100] | 15.05 | 11.40 – 18.70 | **<0.001** |
| MEDIUM [Oleic Acid] × CONC [100] | 21.7 | 18.05 – 25.35 | **<0.001** |
| MEDIUM [Squalene] × CONC [100] | 41.05 | 37.40 – 44.70 | **<0.001** |
| MEDIUM [Cholesterol] × CONC [10] | 24.46 | 20.70 – 28.22 | **<0.001** |
| MEDIUM [Oleic Acid] × CONC [10] | 44.96 | 41.20 – 48.72 | **<0.001** |
| MEDIUM [Squalene] × CONC [10] | 61.93 | 58.05 – 65.80 | **<0.001** |
| MEDIUM [Cholesterol] × CONC [1] | 15.25 | 11.60 – 18.90 | **<0.001** |
| MEDIUM [Oleic Acid] × CONC [1] | 37.3 | 33.65 – 40.95 | **<0.001** |
| MEDIUM [Squalene] × CONC [1] | 53.15 | 49.50 – 56.80 | **<0.001** |
| Observations | 97 | | |
| R^2^ / R^2^ adjusted | 0.987 / 0.984 | | |
