## Supplementary Table 3 for "Skin lipid chemistry influences host-microbiome-pathogen interactions in snake fungal disease (ophidiomycosis)"

**Table S3:** Pairwise contrasts accounting for significant interactions in the linear regression model from Table S3. Estimates shown are *Ophidiomyces ophidiicola* diameter (mm).

| Concentration | Medium | Contrast | Estimate | SE | df | t.ratio | p |
| --- | --- | --- | --- | --- | --- | --- | --- |
| 10,000 | . | Keratin - Cholesterol | 19.200 | 1.3 | 77 | 14.81 | <.001 |
| 10,000 | . | Keratin - Oleic Acid | 44.900 | 1.3 | 77 | 34.64 | <.001 |
| 10,000 | . | Keratin - Squalene | 62.050 | 1.3 | 77 | 47.88 | <.001 |
| 10,000 | . | Cholesterol - Oleic Acid | 25.700 | 1.3 | 77 | 19.83 | <.001 |
| 10,000 | . | Cholesterol - Squalene | 42.850 | 1.3 | 77 | 33.06 | <.001 |
| 10,000 | . | Oleic Acid - Squalene | 17.150 | 1.3 | 77 | 13.23 | <.001 |
| 1,000 | . | Keratin - Cholesterol | 14.950 | 1.3 | 77 | 11.54 | <.001 |
| 1,000 | . | Keratin - Oleic Acid | 36.150 | 1.3 | 77 | 27.89 | <.001 |
| 1,000 | . | Keratin - Squalene | 55.800 | 1.37 | 77 | 40.59 | <.001 |
| 1,000 | . | Cholesterol - Oleic Acid | 21.200 | 1.3 | 77 | 16.36 | <.001 |
| 1,000 | . | Cholesterol - Squalene | 40.850 | 1.37 | 77 | 29.72 | <.001 |
| 1,000 | . | Oleic Acid - Squalene | 19.650 | 1.37 | 77 | 14.29 | <.001 |
| 100 | . | Keratin - Cholesterol | 4.150 | 1.3 | 77 | 3.20 | 0.139 |
| 100 | . | Keratin - Oleic Acid | 23.200 | 1.3 | 77 | 17.90 | <.001 |
| 100 | . | Keratin - Squalene | 21.000 | 1.3 | 77 | 16.20 | <.001 |
| 100 | . | Cholesterol - Oleic Acid | 19.050 | 1.3 | 77 | 14.70 | <.001 |
| 100 | . | Cholesterol - Squalene | 16.850 | 1.3 | 77 | 13.00 | <.001 |
| 100 | . | Oleic Acid - Squalene | -2.200 | 1.3 | 77 | -1.70 | 1 |
| 10 | . | Keratin - Cholesterol | -5.263 | 1.37 | 77 | -3.83 | 0.018 |
| 10 | . | Keratin - Oleic Acid | -0.063 | 1.37 | 77 | -0.05 | 1 |
| 10 | . | Keratin - Squalene | 0.125 | 1.45 | 77 | 0.09 | 1 |
| 10 | . | Cholesterol - Oleic Acid | 5.200 | 1.3 | 77 | 4.01 | 0.010 |
| 10 | . | Cholesterol - Squalene | 5.388 | 1.37 | 77 | 3.92 | 0.013 |
| 10 | . | Oleic Acid - Squalene | 0.188 | 1.37 | 77 | 0.14 | 1 |
| 1 | . | Keratin - Cholesterol | 3.950 | 1.3 | 77 | 3.05 | 0.221 |
| 1 | . | Keratin - Oleic Acid | 7.600 | 1.3 | 77 | 5.86 | <.001 |
| 1 | . | Keratin - Squalene | 8.900 | 1.3 | 77 | 6.87 | <.001 |
| 1 | . | Cholesterol - Oleic Acid | 3.650 | 1.3 | 77 | 2.82 | 0.432 |
| 1 | . | Cholesterol - Squalene | 4.950 | 1.3 | 77 | 3.82 | 0.019 |
| 1 | . | Oleic Acid - Squalene | 1.300 | 1.3 | 77 | 1.00 | 1 |
| . | Keratin | 10,000 - 1,000 | 6.250 | 1.3 | 77 | 4.82 | 0.001 |
| . | Keratin | 10,000 - 100 | 10.050 | 1.3 | 77 | 7.75 | <.001 |
| . | Keratin | 10,000 - 10 | 22.363 | 1.37 | 77 | 16.27 | <.001 |
| . | Keratin | 10,000 - 1 | 14.200 | 1.3 | 77 | 10.96 | <.001 |
| . | Keratin | 1,000 - 100 | 3.800 | 1.3 | 77 | 2.93 | 0.310 |
| . | Keratin | 1,000 - 10 | 16.113 | 1.37 | 77 | 11.72 | <.001 |
| . | Keratin | 1,000 - 1 | 7.950 | 1.3 | 77 | 6.13 | <.001 |
| . | Keratin | 100 - 10 | 12.313 | 1.37 | 77 | 8.96 | <.001 |
| . | Keratin | 100 - 1 | 4.150 | 1.3 | 77 | 3.20 | 0.139 |
| . | Keratin | 10 - 1 | -8.163 | 1.37 | 77 | -5.94 | <.001 |
| . | Cholesterol | 10,000 - 1,000 | 2.000 | 1.3 | 77 | 1.54 | 1 |
| . | Cholesterol | 10,000 - 100 | -5.000 | 1.3 | 77 | -3.86 | 0.017 |
| . | Cholesterol | 10,000 - 10 | -2.100 | 1.3 | 77 | -1.62 | 1 |
| . | Cholesterol | 10,000 - 1 | -1.050 | 1.3 | 77 | -0.81 | 1 |
| . | Cholesterol | 1,000 - 100 | -7.000 | 1.3 | 77 | -5.40 | <.001 |
| . | Cholesterol | 1,000 - 10 | -4.100 | 1.3 | 77 | -3.16 | 0.156 |
| . | Cholesterol | 1,000 - 1 | -3.050 | 1.3 | 77 | -2.35 | 1 |
| . | Cholesterol | 100 - 10 | 2.900 | 1.3 | 77 | 2.24 | 1 |
| . | Cholesterol | 100 - 1 | 3.950 | 1.3 | 77 | 3.05 | 0.221 |
| . | Cholesterol | 10 - 1 | 1.050 | 1.3 | 77 | 0.81 | 1 |
| . | Oleic Acid | 10,000 - 1,000 | -2.500 | 1.3 | 77 | -1.93 | 1 |
| . | Oleic Acid | 10,000 - 100 | -11.650 | 1.3 | 77 | -8.99 | <.001 |
| . | Oleic Acid | 10,000 - 10 | -22.600 | 1.3 | 77 | -17.44 | <.001 |
| . | Oleic Acid | 10,000 - 1 | -23.100 | 1.3 | 77 | -17.82 | <.001 |
| . | Oleic Acid | 1,000 - 100 | -9.150 | 1.3 | 77 | -7.06 | <.001 |
| . | Oleic Acid | 1,000 - 10 | -20.100 | 1.3 | 77 | -15.51 | <.001 |
| . | Oleic Acid | 1,000 - 1 | -20.600 | 1.3 | 77 | -15.89 | <.001 |
| . | Oleic Acid | 100 - 10 | -10.950 | 1.3 | 77 | -8.45 | <.001 |
| . | Oleic Acid | 100 - 1 | -11.450 | 1.3 | 77 | -8.83 | <.001 |
| . | Oleic Acid | 10 - 1 | -0.500 | 1.3 | 77 | -0.39 | 1 |
| . | Squalene | 10,000 - 1,000 | 0.000 | 1.37 | 77 | 0.00 | 1 |
| . | Squalene | 10,000 - 100 | -31.000 | 1.3 | 77 | -23.92 | <.001 |
| . | Squalene | 10,000 - 10 | -39.563 | 1.37 | 77 | -28.78 | <.001 |
| . | Squalene | 10,000 - 1 | -38.950 | 1.3 | 77 | -30.05 | <.001 |
| . | Squalene | 1,000 - 100 | -31.000 | 1.37 | 77 | -22.55 | <.001 |
| . | Squalene | 1,000 - 10 | -39.563 | 1.45 | 77 | -27.30 | <.001 |
| . | Squalene | 1,000 - 1 | -38.950 | 1.37 | 77 | -28.33 | <.001 |
| . | Squalene | 100 - 10 | -8.563 | 1.37 | 77 | -6.23 | <.001 |
| . | Squalene | 100 - 1 | -7.950 | 1.3 | 77 | -6.13 | <.001 |
| . | Squalene | 10 - 1 | 0.613 | 1.37 | 77 | 0.45 | 1 |
