## Supplementary Table 4 for "Skin lipid chemistry influences host-microbiome-pathogen interactions in snake fungal disease (ophidiomycosis)"

**Table S4:** Linear regression models examining the diameter of *Ophidiomyces ophidiicola* growth (mm) within each of the four minimal media treatments relative to M9 minimal media controls. Variables shown represent the medium and its concentration (i.e., Keratin 10,000 represents Keratin at 10,000 ppm).

|  | Estimate | Std. Error | t value | p |
| --- | --- | --- | --- | --- |
| (Intercept) | 41.25 | 0.6319 | 65.277 | < 0.0001 |
| Keratin 10,000 | 20.8 | 1.3255 | 15.692 | < 0.0001 |
| Keratin 1,000 | 14.55 | 1.3255 | 10.977 | < 0.0001 |
| Keratin 100 | 10.75 | 1.3255 | 8.11 | < 0.0001 |
| Keratin 10 | -1.5625 | 1.4479 | -1.079 | 0.288 |
| Keratin 1 | 6.6 | 1.3255 | 4.979 | < 0.0001 |
| (Intercept) | 41.25 | 0.8864 | 46.537 | < 0.0001 |
| Cholesterol 10,000 | 1.6 | 1.8593 | 0.861 | 0.3952 |
| Cholesterol 1,000 | -0.4 | 1.8593 | -0.215 | 0.8309 |
| Cholesterol 100 | 6.6 | 1.8593 | 3.55 | 0.0011 |
| Cholesterol 10 | 3.7 | 1.8593 | 1.99 | 0.0542 |
| Cholesterol 1 | 2.65 | 1.8593 | 1.425 | 0.1627 |
| (Intercept) | 41.25 | 0.6549 | 62.991 | < 0.0001 |
| Oleic Acid 10,000 | -24.1 | 1.3736 | -17.55 | < 0.0001 |
| Oleic Acid 1,000 | -21.6 | 1.3736 | -15.73 | < 0.0001 |
| Oleic Acid 100 | -12.45 | 1.3736 | -9.063 | < 0.0001 |
| Oleic Acid 10 | -1.5 | 1.3736 | -1.092 | 0.282 |
| Oleic Acid 1 | -1 | 1.3736 | -0.728 | 0.471 |
| (Intercept) | 41.25 | 0.6437 | 64.086 | < 0.0001 |
| Squalene 10,000 | -41.25 | 1.3502 | -30.55 | < 0.0001 |
| Squalene 1,000 | -41.25 | 1.4748 | -27.97 | < 0.0001 |
| Squalene 100 | -10.25 | 1.3502 | -7.592 | < 0.0001 |
| Squalene 10 | -1.6875 | 1.4748 | -1.144 | 0.2605 |
| Squalene 1 | -2.3 | 1.3502 | -1.703 | 0.0976 |
