## Supplementary Table 5 for "Skin lipid chemistry influences host-microbiome-pathogen interactions in snake fungal disease (ophidiomycosis)"

**Table S5:** Linear regression models examining the diameter of *Ophidiomyces ophidiicola* growth (mm) across mixed media plates with 10,000 ppm keratin as the reference level.

|  | Estimate | Std. Error | t value | Pr(>\|t\|) |
| --- | --- | --- | --- | --- |
| (Intercept) | 62.05 | 0.2994 | 207.217 | <0.0001 |
| Keratin+Cholesterol | 0.95 | 0.4492 | 2.115 | 0.0516 |
| Keratin+Squalene | -62.05 | 0.4235 | -146.525 | <0.0001 |
| Keratin+Oleic Acid | -28.25 | 0.4235 | -66.71 | <0.0001 |
