## Supplementary Table 6 for "Skin lipid chemistry influences host-microbiome-pathogen interactions in snake fungal disease (ophidiomycosis)"

**Supplementary Table 6:** Growth of bacterial species (isolate ID given in parentheses), in optical density, in the presence and absence of *O. ophidiicola* (*Oo*) grown in co-culture using an ANOVA. Fungal growth in the presence of each bacterial isolate (in unit) on keratin media across time using a linear fixed effects model.

| Bacterial species | Effect of *Oo* | | Effect of media | | Fungal growth | |
| --- | --- | --- | --- | --- | --- | --- |
|  | F value | *P* value | F value | *P* value | β ± se | *P* value |
| *Acinetobacter guillouiae* (BR1.9) | 3.27 | 0.11 | 1.35 | 0.28 | 0.02 ± 0.01 | 0.09 |
| *Chryseobacterium* sp. (BR1.10) | 5.62 | **0.045** | 34.2 | **<0.001** | -0.09 ± 0.01 | **<0.001** |
| *Flavobacterium odoratimimum* (TR087-7.2) | 0.27 | 0.62 | 4.83 | 0.06 | -0.03 ± 0.01 | **0.006** |
| *Lelliottia nimipressuralis* (BR1.11) | 0.01 | 0.94 | 19.0 | **0.002** | -0.07 ± 0.01 | **<0.001** |
| *Mammaliicoccus sciuri* (BR1.7) | 5.49 | **0.047** | 4.94 | 0.057 | -0.02 ± 0.01 | 0.19 |
| *Stenotrophomonas* sp. (EKS20.5) | 0.70 | 0.43 | 755 | **<0.001** | -0.10 ± 0.02 | **<0.001** |
