## Supplementary Table 7 for "Skin lipid chemistry influences host-microbiome-pathogen interactions in snake fungal disease (ophidiomycosis)"

**Supplementary Table 7:** Pairwise contrasts using Tukey’s HSD of bacterial growth across the different media types in the fungal spent media experiment.

| Isolate | Bacterial Species | Group1 | Group2 | df | Statistic | p |
| --- | --- | --- | --- | --- | --- | --- |
| blank |  | m9 | keratin | 85 | 4.465 | <0.001 |
| blank |  | m9 | 5d | 85 | 4.024 | <0.001 |
| blank |  | m9 | 9d | 85 | 4.485 | <0.001 |
| blank |  | m9 | 13d | 85 | 3.526 | <0.001 |
| blank |  | keratin | 5d | 85 | -0.441 | 0.660 |
| blank |  | keratin | 9d | 85 | 0.020 | 0.984 |
| blank |  | keratin | 13d | 85 | -0.939 | 0.350 |
| blank |  | 5d | 9d | 85 | 0.461 | 0.646 |
| blank |  | 5d | 13d | 85 | -0.498 | 0.620 |
| blank |  | 9d | 13d | 85 | -0.959 | 0.340 |
| EKS20.5 | *Stenotrophomonas maltophilia* | m9 | keratin | 10 | -9.628 | <0.001 |
| EKS20.5 | *Stenotrophomonas maltophilia* | m9 | 5d | 10 | -38.734 | <0.001 |
| EKS20.5 | *Stenotrophomonas maltophilia* | m9 | 9d | 10 | -24.948 | <0.001 |
| EKS20.5 | *Stenotrophomonas maltophilia* | m9 | 13d | 10 | -20.841 | <0.001 |
| EKS20.5 | *Stenotrophomonas maltophilia* | keratin | 5d | 10 | -29.106 | <0.001 |
| EKS20.5 | *Stenotrophomonas maltophilia* | keratin | 9d | 10 | -15.321 | <0.001 |
| EKS20.5 | *Stenotrophomonas maltophilia* | keratin | 13d | 10 | -11.213 | <0.001 |
| EKS20.5 | *Stenotrophomonas maltophilia* | 5d | 9d | 10 | 13.785 | <0.001 |
| EKS20.5 | *Stenotrophomonas maltophilia* | 5d | 13d | 10 | 17.893 | <0.001 |
| EKS20.5 | *Stenotrophomonas maltophilia* | 9d | 13d | 10 | 4.108 | 0.002 |
| BR1.9 | *Acinetobacter guillouiae* | m9 | keratin | 10 | -1.178 | 0.266 |
| BR1.9 | *Acinetobacter guillouiae* | m9 | 5d | 10 | -1.056 | 0.316 |
| BR1.9 | *Acinetobacter guillouiae* | m9 | 9d | 10 | -4.137 | 0.002 |
| BR1.9 | *Acinetobacter guillouiae* | m9 | 13d | 10 | 0.118 | 0.909 |
| BR1.9 | *Acinetobacter guillouiae* | keratin | 5d | 10 | 0.122 | 0.905 |
| BR1.9 | *Acinetobacter guillouiae* | keratin | 9d | 10 | -2.959 | 0.014 |
| BR1.9 | *Acinetobacter guillouiae* | keratin | 13d | 10 | 1.296 | 0.224 |
| BR1.9 | *Acinetobacter guillouiae* | 5d | 9d | 10 | -3.081 | 0.012 |
| BR1.9 | *Acinetobacter guillouiae* | 5d | 13d | 10 | 1.174 | 0.268 |
| BR1.9 | *Acinetobacter guillouiae* | 9d | 13d | 10 | 4.255 | 0.002 |
| BR1.1 | *Chryseobacterium* sp. | m9 | keratin | 10 | 1.886 | 0.089 |
| BR1.1 | *Chryseobacterium* sp. | m9 | 5d | 10 | -15.185 | <0.001 |
| BR1.1 | *Chryseobacterium* sp. | m9 | 9d | 10 | -1.530 | 0.157 |
| BR1.1 | *Chryseobacterium* sp. | m9 | 13d | 10 | 0.078 | 0.939 |
| BR1.1 | *Chryseobacterium* sp. | keratin | 5d | 10 | -17.071 | <0.001 |
| BR1.1 | *Chryseobacterium* sp. | keratin | 9d | 10 | -3.417 | 0.007 |
| BR1.1 | *Chryseobacterium* sp. | keratin | 13d | 10 | -1.808 | 0.101 |
| BR1.1 | *Chryseobacterium* sp. | 5d | 9d | 10 | 13.654 | <0.001 |
| BR1.1 | *Chryseobacterium* sp. | 5d | 13d | 10 | 15.263 | <0.001 |
| BR1.1 | *Chryseobacterium* sp. | 9d | 13d | 10 | 1.609 | 0.139 |
| BR1.10 | *Chryseobacterium* sp. | m9 | keratin | 10 | 4.261 | 0.002 |
| BR1.10 | *Chryseobacterium* sp. | m9 | 5d | 10 | -29.630 | <0.001 |
| BR1.10 | *Chryseobacterium* sp. | m9 | 9d | 10 | -2.925 | 0.015 |
| BR1.10 | *Chryseobacterium* sp. | m9 | 13d | 10 | -2.399 | 0.037 |
| BR1.10 | *Chryseobacterium* sp. | keratin | 5d | 10 | -33.890 | <0.001 |
| BR1.10 | *Chryseobacterium* sp. | keratin | 9d | 10 | -7.186 | <0.001 |
| BR1.10 | *Chryseobacterium* sp. | keratin | 13d | 10 | -6.660 | <0.001 |
| BR1.10 | *Chryseobacterium* sp. | 5d | 9d | 10 | 26.705 | <0.001 |
| BR1.10 | *Chryseobacterium* sp. | 5d | 13d | 10 | 27.230 | <0.001 |
| BR1.10 | *Chryseobacterium* sp. | 9d | 13d | 10 | 0.526 | 0.611 |
| BR1.6 | *Mammaliicoccus sciuri* | m9 | keratin | 10 | -2.132 | 0.059 |
| BR1.6 | *Mammaliicoccus sciuri* | m9 | 5d | 10 | -8.590 | <0.001 |
| BR1.6 | *Mammaliicoccus sciuri* | m9 | 9d | 10 | -3.402 | 0.007 |
| BR1.6 | *Mammaliicoccus sciuri* | m9 | 13d | 10 | -11.391 | <0.001 |
| BR1.6 | *Mammaliicoccus sciuri* | keratin | 5d | 10 | -6.458 | <0.001 |
| BR1.6 | *Mammaliicoccus sciuri* | keratin | 9d | 10 | -1.271 | 0.233 |
| BR1.6 | *Mammaliicoccus sciuri* | keratin | 13d | 10 | -9.259 | <0.001 |
| BR1.6 | *Mammaliicoccus sciuri* | 5d | 9d | 10 | 5.187 | <0.001 |
| BR1.6 | *Mammaliicoccus sciuri* | 5d | 13d | 10 | -2.801 | 0.019 |
| BR1.6 | *Mammaliicoccus sciuri* | 9d | 13d | 10 | -7.988 | <0.001 |
| EKS17.1 | *Sphingomonas* sp. | m9 | keratin | 10 | 0.115 | 0.910 |
| EKS17.1 | *Sphingomonas* sp. | m9 | 5d | 10 | 0.224 | 0.827 |
| EKS17.1 | *Sphingomonas* sp. | m9 | 9d | 10 | 0.860 | 0.410 |
| EKS17.1 | *Sphingomonas* sp. | m9 | 13d | 10 | 1.215 | 0.252 |
| EKS17.1 | *Sphingomonas* sp. | keratin | 5d | 10 | 0.109 | 0.915 |
| EKS17.1 | *Sphingomonas* sp. | keratin | 9d | 10 | 0.745 | 0.474 |
| EKS17.1 | *Sphingomonas* sp. | keratin | 13d | 10 | 1.100 | 0.297 |
| EKS17.1 | *Sphingomonas* sp. | 5d | 9d | 10 | 0.636 | 0.539 |
| EKS17.1 | *Sphingomonas* sp. | 5d | 13d | 10 | 0.991 | 0.345 |
| EKS17.1 | *Sphingomonas* sp. | 9d | 13d | 10 | 0.355 | 0.730 |
| TR087-6.2 |  | m9 | keratin | 10 | -0.416 | 0.686 |
| TR087-6.2 |  | m9 | 5d | 10 | -15.397 | <0.001 |
| TR087-6.2 |  | m9 | 9d | 10 | -3.067 | 0.012 |
| TR087-6.2 |  | m9 | 13d | 10 | -9.202 | <0.001 |
| TR087-6.2 |  | keratin | 5d | 10 | -14.982 | <0.001 |
| TR087-6.2 |  | keratin | 9d | 10 | -2.651 | 0.024 |
| TR087-6.2 |  | keratin | 13d | 10 | -8.786 | <0.001 |
| TR087-6.2 |  | 5d | 9d | 10 | 12.331 | <0.001 |
| TR087-6.2 |  | 5d | 13d | 10 | 6.196 | <0.001 |
| TR087-6.2 |  | 9d | 13d | 10 | -6.135 | <0.001 |
| EKS20.21 |  | m9 | keratin | 10 | -0.353 | 0.731 |
| EKS20.21 |  | m9 | 5d | 10 | -27.638 | <0.001 |
| EKS20.21 |  | m9 | 9d | 10 | -15.226 | <0.001 |
| EKS20.21 |  | m9 | 13d | 10 | -17.243 | <0.001 |
| EKS20.21 |  | keratin | 5d | 10 | -27.284 | <0.001 |
| EKS20.21 |  | keratin | 9d | 10 | -14.873 | <0.001 |
| EKS20.21 |  | keratin | 13d | 10 | -16.890 | <0.001 |
| EKS20.21 |  | 5d | 9d | 10 | 12.412 | <0.001 |
| EKS20.21 |  | 5d | 13d | 10 | 10.395 | <0.001 |
| EKS20.21 |  | 9d | 13d | 10 | -2.017 | 0.071 |
| BR1.7 | *Mammaliicoccus sciuri* | m9 | keratin | 10 | 1.297 | 0.224 |
| BR1.7 | *Mammaliicoccus sciuri* | m9 | 5d | 10 | -11.877 | <0.001 |
| BR1.7 | *Mammaliicoccus sciuri* | m9 | 9d | 10 | -0.763 | 0.463 |
| BR1.7 | *Mammaliicoccus sciuri* | m9 | 13d | 10 | -5.706 | <0.001 |
| BR1.7 | *Mammaliicoccus sciuri* | keratin | 5d | 10 | -13.173 | <0.001 |
| BR1.7 | *Mammaliicoccus sciuri* | keratin | 9d | 10 | -2.060 | 0.066 |
| BR1.7 | *Mammaliicoccus sciuri* | keratin | 13d | 10 | -7.003 | <0.001 |
| BR1.7 | *Mammaliicoccus sciuri* | 5d | 9d | 10 | 11.113 | <0.001 |
| BR1.7 | *Mammaliicoccus sciuri* | 5d | 13d | 10 | 6.170 | <0.001 |
| BR1.7 | *Mammaliicoccus sciuri* | 9d | 13d | 10 | -4.943 | <0.001 |
| EKS17.21 |  | m9 | keratin | 10 | 0.901 | 0.389 |
| EKS17.21 |  | m9 | 5d | 10 | 1.129 | 0.285 |
| EKS17.21 |  | m9 | 9d | 10 | -0.081 | 0.937 |
| EKS17.21 |  | m9 | 13d | 10 | 0.706 | 0.496 |
| EKS17.21 |  | keratin | 5d | 10 | 0.228 | 0.824 |
| EKS17.21 |  | keratin | 9d | 10 | -0.982 | 0.349 |
| EKS17.21 |  | keratin | 13d | 10 | -0.195 | 0.849 |
| EKS17.21 |  | 5d | 9d | 10 | -1.210 | 0.254 |
| EKS17.21 |  | 5d | 13d | 10 | -0.423 | 0.681 |
| EKS17.21 |  | 9d | 13d | 10 | 0.787 | 0.449 |
| TR087-7.2 | *Flavobacterium odoratimimum* | m9 | keratin | 10 | -0.436 | 0.672 |
| TR087-7.2 | *Flavobacterium odoratimimum* | m9 | 5d | 10 | -21.376 | <0.001 |
| TR087-7.2 | *Flavobacterium odoratimimum* | m9 | 9d | 10 | -23.617 | <0.001 |
| TR087-7.2 | *Flavobacterium odoratimimum* | m9 | 13d | 10 | -21.723 | <0.001 |
| TR087-7.2 | *Flavobacterium odoratimimum* | keratin | 5d | 10 | -20.939 | <0.001 |
| TR087-7.2 | *Flavobacterium odoratimimum* | keratin | 9d | 10 | -23.180 | <0.001 |
| TR087-7.2 | *Flavobacterium odoratimimum* | keratin | 13d | 10 | -21.287 | <0.001 |
| TR087-7.2 | *Flavobacterium odoratimimum* | 5d | 9d | 10 | -2.241 | 0.049 |
| TR087-7.2 | *Flavobacterium odoratimimum* | 5d | 13d | 10 | -0.347 | 0.735 |
| TR087-7.2 | *Flavobacterium odoratimimum* | 9d | 13d | 10 | 1.894 | 0.088 |
| EKS20.8 | *Paenarthrobacter* sp. | m9 | keratin | 10 | 2.390 | 0.038 |
| EKS20.8 | *Paenarthrobacter* sp. | m9 | 5d | 10 | -5.457 | <0.001 |
| EKS20.8 | *Paenarthrobacter* sp. | m9 | 9d | 10 | 1.323 | 0.215 |
| EKS20.8 | *Paenarthrobacter* sp. | m9 | 13d | 10 | -8.079 | <0.001 |
| EKS20.8 | *Paenarthrobacter* sp. | keratin | 5d | 10 | -7.847 | <0.001 |
| EKS20.8 | *Paenarthrobacter* sp. | keratin | 9d | 10 | -1.067 | 0.311 |
| EKS20.8 | *Paenarthrobacter* sp. | keratin | 13d | 10 | -10.469 | <0.001 |
| EKS20.8 | *Paenarthrobacter* sp. | 5d | 9d | 10 | 6.780 | <0.001 |
| EKS20.8 | *Paenarthrobacter* sp. | 5d | 13d | 10 | -2.622 | 0.026 |
| EKS20.8 | *Paenarthrobacter* sp. | 9d | 13d | 10 | -9.402 | <0.001 |
| EKS17.8 | *Stenotrophomonas* sp. | m9 | keratin | 10 | -3.971 | 0.003 |
| EKS17.8 | *Stenotrophomonas* sp. | m9 | 5d | 10 | -39.566 | <0.001 |
| EKS17.8 | *Stenotrophomonas* sp. | m9 | 9d | 10 | -21.498 | <0.001 |
| EKS17.8 | *Stenotrophomonas* sp. | m9 | 13d | 10 | -17.766 | <0.001 |
| EKS17.8 | *Stenotrophomonas* sp. | keratin | 5d | 10 | -35.594 | <0.001 |
| EKS17.8 | *Stenotrophomonas* sp. | keratin | 9d | 10 | -17.527 | <0.001 |
| EKS17.8 | *Stenotrophomonas* sp. | keratin | 13d | 10 | -13.795 | <0.001 |
| EKS17.8 | *Stenotrophomonas* sp. | 5d | 9d | 10 | 18.067 | <0.001 |
| EKS17.8 | *Stenotrophomonas* sp. | 5d | 13d | 10 | 21.799 | <0.001 |
| EKS17.8 | *Stenotrophomonas* sp. | 9d | 13d | 10 | 3.732 | 0.004 |
| BR1.11 | *Leilliottia nimipressularis* | m9 | keratin | 10 | 0.172 | 0.867 |
| BR1.11 | *Leilliottia nimipressularis* | m9 | 5d | 10 | -43.183 | <0.001 |
| BR1.11 | *Leilliottia nimipressularis* | m9 | 9d | 10 | -36.852 | <0.001 |
| BR1.11 | *Leilliottia nimipressularis* | m9 | 13d | 10 | -29.178 | <0.001 |
| BR1.11 | *Leilliottia nimipressularis* | keratin | 5d | 10 | -43.355 | <0.001 |
| BR1.11 | *Leilliottia nimipressularis* | keratin | 9d | 10 | -37.024 | <0.001 |
| BR1.11 | *Leilliottia nimipressularis* | keratin | 13d | 10 | -29.350 | <0.001 |
| BR1.11 | *Leilliottia nimipressularis* | 5d | 9d | 10 | 6.331 | <0.001 |
| BR1.11 | *Leilliottia nimipressularis* | 5d | 13d | 10 | 14.005 | <0.001 |
| BR1.11 | *Leilliottia nimipressularis* | 9d | 13d | 10 | 7.674 | <0.001 |
| TR087-5.1 | *Aeromonas hydrophila* | m9 | keratin | 10 | -2.585 | 0.027 |
| TR087-5.1 | *Aeromonas hydrophila* | m9 | 5d | 10 | -8.068 | <0.001 |
| TR087-5.1 | *Aeromonas hydrophila* | m9 | 9d | 10 | -5.061 | <0.001 |
| TR087-5.1 | *Aeromonas hydrophila* | m9 | 13d | 10 | -3.218 | 0.009 |
| TR087-5.1 | *Aeromonas hydrophila* | keratin | 5d | 10 | -5.484 | <0.001 |
| TR087-5.1 | *Aeromonas hydrophila* | keratin | 9d | 10 | -2.477 | 0.033 |
| TR087-5.1 | *Aeromonas hydrophila* | keratin | 13d | 10 | -0.634 | 0.540 |
| TR087-5.1 | *Aeromonas hydrophila* | 5d | 9d | 10 | 3.007 | 0.013 |
| TR087-5.1 | *Aeromonas hydrophila* | 5d | 13d | 10 | 4.850 | <0.001 |
| TR087-5.1 | *Aeromonas hydrophila* | 9d | 13d | 10 | 1.843 | 0.095 |
| EKS20.10 | *Stenotrophomonas* sp. | m9 | keratin | 10 | 1.322 | 0.215 |
| EKS20.10 | *Stenotrophomonas* sp. | m9 | 5d | 10 | -24.460 | <0.001 |
| EKS20.10 | *Stenotrophomonas* sp. | m9 | 9d | 10 | -15.387 | <0.001 |
| EKS20.10 | *Stenotrophomonas* sp. | m9 | 13d | 10 | -14.826 | <0.001 |
| EKS20.10 | *Stenotrophomonas* sp. | keratin | 5d | 10 | -25.782 | <0.001 |
| EKS20.10 | *Stenotrophomonas* sp. | keratin | 9d | 10 | -16.709 | <0.001 |
| EKS20.10 | *Stenotrophomonas* sp. | keratin | 13d | 10 | -16.149 | <0.001 |
| EKS20.10 | *Stenotrophomonas* sp. | 5d | 9d | 10 | 9.073 | <0.001 |
| EKS20.10 | *Stenotrophomonas* sp. | 5d | 13d | 10 | 9.633 | <0.001 |
| EKS20.10 | *Stenotrophomonas* sp. | 9d | 13d | 10 | 0.560 | 0.588 |
| BR1.2 | *Acinetobacter dispersus* | m9 | keratin | 10 | 4.060 | 0.002 |
| BR1.2 | *Acinetobacter dispersus* | m9 | 5d | 10 | -28.377 | <0.001 |
| BR1.2 | *Acinetobacter dispersus* | m9 | 9d | 10 | -28.136 | <0.001 |
| BR1.2 | *Acinetobacter dispersus* | m9 | 13d | 10 | -8.016 | <0.001 |
| BR1.2 | *Acinetobacter dispersus* | keratin | 5d | 10 | -32.437 | <0.001 |
| BR1.2 | *Acinetobacter dispersus* | keratin | 9d | 10 | -32.195 | <0.001 |
| BR1.2 | *Acinetobacter dispersus* | keratin | 13d | 10 | -12.076 | <0.001 |
| BR1.2 | *Acinetobacter dispersus* | 5d | 9d | 10 | 0.242 | 0.814 |
| BR1.2 | *Acinetobacter dispersus* | 5d | 13d | 10 | 20.361 | <0.001 |
| BR1.2 | *Acinetobacter dispersus* | 9d | 13d | 10 | 20.119 | <0.001 |
| EKS17.29 | *Streptomyces* sp. | m9 | keratin | 10 | 6.390 | <0.001 |
| EKS17.29 | *Streptomyces* sp. | m9 | 5d | 10 | 3.225 | 0.009 |
| EKS17.29 | *Streptomyces* sp. | m9 | 9d | 10 | 4.450 | 0.001 |
| EKS17.29 | *Streptomyces* sp. | m9 | 13d | 10 | 4.627 | <0.001 |
| EKS17.29 | *Streptomyces* sp. | keratin | 5d | 10 | -3.165 | 0.010 |
| EKS17.29 | *Streptomyces* sp. | keratin | 9d | 10 | -1.939 | 0.081 |
| EKS17.29 | *Streptomyces* sp. | keratin | 13d | 10 | -1.763 | 0.108 |
| EKS17.29 | *Streptomyces* sp. | 5d | 9d | 10 | 1.225 | 0.249 |
| EKS17.29 | *Streptomyces* sp. | 5d | 13d | 10 | 1.402 | 0.191 |
| EKS17.29 | *Streptomyces* sp. | 9d | 13d | 10 | 0.177 | 0.863 |
