## Supplementary Table 8 for "Skin lipid chemistry influences host-microbiome-pathogen interactions in snake fungal disease (ophidiomycosis)"

**Supplementary Table 8:** Putative biosynthetic gene clusters from 26 bacterial isolates obtained from free-ranging snakes. Inhibits *Oo* status represents *in vitro* assay results from Hill et al. ^60^.

| Isolate id | Host species | gtdb_species | Inhibits *Oo* | Host SFD status | total bgc regions | Non saccharide, fatty acid, and saccharide-fatty acid regions | saccharide, fatty acid, and saccharide-fatty acid regions | antiSMASH siderophore BGC regions | Notable rediscoveries |
| --- | --- | --- | --- | --- | --- | --- | --- | --- | --- |
| BR 1.10 | *Coluber constrictor* | s__Chryseobacterium sp024158915 | 0 | 1 | 32 (1) | 8 (1) | 24 (0) | 1 | flexirubin |
| BR 1.11 | *Coluber constrictor* | s__Lelliottia nimipressuralis | 1 | 1 | 17 (2) | 4 (1) | 13 (1) | 0 | arylpolyene |
| BR 1.12 | *Coluber constrictor* | s__Acinetobacter guillouiae | 0 | 1 | 9 (3) | 4 (2) | 5 (1) | 2 | acinetoferrin |
| BR 1.2 | *Coluber constrictor* | s__Acinetobacter dispersus | 0 | 1 | 13 (1) | 8 (1) | 5 (0) | 1 | - |
| BR 1.4 | *Coluber constrictor* | s__Empedobacter tilapiae | 0 | 1 | 22 (6) | 1 (0) | 21 (6) | 0 | flexirubin |
| BR 1.5 | *Coluber constrictor* | s__Flavobacterium odoratum_A | 0 | 1 | 22 (9) | 5 (1) | 17 (8) | 0 | flexirubin |
| BR 1.6 | *Coluber constrictor* | s__Mammaliicoccus sciuri | 0 | 1 | 11 (2) | 5 (2) | 6 (0) | 1 | - |
| BR 1.7 | *Coluber constrictor* | s__Mammaliicoccus sciuri | 0 | 1 | 13 (2) | 6 (2) | 7 (0) | 1 | - |
| BR 1.8 | *Coluber constrictor* | s__Acinetobacter guillouiae | 0 | 1 | 11 (4) | 5 (2) | 6 (2) | 1 | - |
| BR 1.9 | *Coluber constrictor* | s__Acinetobacter guillouiae | 0 | 1 | 11 (4) | 5 (2) | 6 (2) | 1 | - |
| EKS17.1 | *Pantherophis spiloides* | s__Sphingomonas sp. | NA | NA | 29 (9) | 13 (3) | 16 (6) | 0 | - |
| EKS17.15 | *Pantherophis spiloides* | g__Pristimantibacillus | NA | NA | 25 (11) | 6 (3) | 19 (8) | 0 | paeninodin |
| EKS17.19 | *Pantherophis spiloides* | g__Microbacterium | NA | NA | 18 (4) | 7 (2) | 11 (2) | 0 | - |
| EKS17.29 | *Pantherophis spiloides* | g__Streptomyces | NA | NA | 74 (50) | 52 (36) | 22 (14) | 2 | geosmin, SGR PTMs, desferrioxamin B, endophenazine, keywimycin, ectoine, streptobactin, coelichelin, melanin, AmfS, streptothricin |
| EKS20.1 | *Lampropeltis nigra* | f__Dermatophilaceae | NA | NA | 18 (7) | 3 (1) | 15 (6) | 1 | desferrioxamin B |
| EKS20.3 | *Lampropeltis nigra* | g__Lysinibacillus | NA | NA | 22 (9) | 14 (8) | 8 (1) | 1 | - |
| EKS20.4 | *Lampropeltis nigra* | s__Rhodococcus erythropolis_D | NA | NA | 34 (11) | 18 (7) | 16 (4) | 0 | heterobactin, ectoine |
| EKS20.8 | *Lampropeltis nigra* | s__Paenarthrobacter sp900106835 | NA | NA | 26 (6) | 9 (1) | 17 (5) | 1 | desferrioxamine E |
| TR087-5.1 | *Crotalus horridus* | s__Aeromonas hydrophila | 1 | 0 | 24 (4) | 5 (1) | 19 (3) | 1 | amonavactin P 750 |
| TR087-5.3 | *Crotalus horridus* | s__Morganella morganii | 1 | 0 | 11 (0) | 3 (0) | 8 (0) | 0 | - |
| TR087-6.1 | *Crotalus horridus* | s__Serratia_J liquefaciens | 0 | 0 | 23 (4) | 8 (4) | 15 (0) | 0 | arylpolyene |
| TR087-7.2 | *Crotalus horridus* | s__Flavobacterium odoratimimum | NA | 0 | 17 (4) | 1 (0) | 16 (4) | 0 | flexirubin |
| TR087-7.4 | *Crotalus horridus* | s__Morganella morganii | 1 | 0 | 11 (0) | 3 (0) | 8 (0) | 0 | - |
| TR087-7.6 | *Crotalus horridus* | s__Morganella morganii | 1 | 0 | 11 (0) | 3 (0) | 8 (0) | 0 | - |
| TR087-6.4s | *Crotalus horridus* | s__Stenotrophomonas maltophilia | 1 | 0 | 17 (0) | 4 (0) | 13 (0) | 0 | arylpolyene |
