## Supplementary Table 9 for "Skin lipid chemistry influences host-microbiome-pathogen interactions in snake fungal disease (ophidiomycosis)"

**Supplementary Table 9:** Diversity of putative biosynthetic gene clusters identified in the *Ophidiomyces ophidiicola* genome.

| **Functional class** | **Total count** | **BGCs** | **Type** | **From** | **To** | **Antismash - most similar known cluster** | **Similarity (%)** |
| --- | --- | --- | --- | --- | --- | --- | --- |
| NRPS | 7 | Region 1.1 | NRPS | 88,084 | 135,606 | Unknown |  |
|  |  | Region 7.1 | NRPS | 110,852 | 175,503 | Unknown |  |
|  |  | Region 10.1 | NRPS | 367,160 | 419,610 | Unknown |  |
|  |  | Region 12.1 | NRPS | 7,513 | 59,539 | Unknown |  |
|  |  | Region 25.1 | NRPS | 106,371 | 151,167 | penigainamide A/penigainamide B/penigainamide C/adametizine A/FA2097/outovirin A/outovirin C/pretrichodermamide C | 65 |
|  |  | Region 28.1 | NRPS | 67,811 | 111,125 | griseofulvin/epidechlorogriseofulvin/norlichexanthone/  dehydrogriseofulvin/4-desmethylgriseofulvin/  griseoxanthone B | 14 |
|  |  | Region 41.1 | NRPS | 29,278 | 84,926 | Unknown |  |
| NRPS-like | 2 | Region 7.2 | NRPS-like | 274,992 | 320,158 | Unknown |  |
|  |  | Region 19.1 | NRPS-like | 214,644 | 259,023 | Unknown |  |
| Hybrids | 3 | Region 2.2 | T1PKS, NRPS | 111,409 | 168,539 | chaetolivacine A/chaetolivacine B/chaetolivacine C | 100 |
|  |  | Region 4.1 | NRPS-like, NRPS | 385,751 | 436,864 | hancockiamide E/hancockiamide B/hancockiamide C/hancockiamide D/hancockiamide A/hancockiamide F | 58 |
|  |  | Region 26.1 | NRPS,indole | 95,892 | 141,908 | okaramine B |  |
| Terpenes | 4 | Region 2.1 | Terpene | 79,007 | 106,571 | clavaric acid | 100 |
|  |  | Region 7.3 | Terpene | 761,013 | 782,311 | Unknown |  |
|  |  | Region 9.1 | Terpene | 533,710 | 556,928 | Unknown |  |
|  |  | Region 10.2 | Terpene | 459,313 | 482,247 | Squalestatin S1 | 40 |
| Polyketides | 4 | Region 2.3 | T1PKS | 306,357 | 353,048 | Unknown |  |
|  |  | Region 22.1 | T1PKS | 83,168 | 145,539 | Unknown |  |
|  |  | Region 37.1 | T1PKS | 110,209 | 158,441 | Unknown |  |
|  |  | Region 48.1 | T1PKS | 54,792 | 89,786 | Unknown |  |
| Indole | 1 | Region 3.1 | Indole | 181,295 | 202,503 | Unknown |  |
| Betalactone | 1 | Region 11.1 | Betalactone | 409,273 | 437,842 | Unknown |  |
| NRP | 1 | Region 39.1 | NRP-metallophore,NRPS | 28,884 | 84,053 | metachelin C/metachelin A/metachelin A-CE/metachelin B/dimerumic acid 11-mannoside/dimerumic acid | 62 |
| RiPP | 1 | Region 5.1 | Fungal-RiPP-like | 715,913 | 819,846 | Unknown |  |
| Isocyanide | 1 | Region 32.1 | Isocyanide | 105676 | 147727 | Unknown |  |
